# Differentiation of *Toxoplasma* into latent forms is linked to central carbon metabolism and requires a GID/CTLH-type E3 ubiquitin ligase

**DOI:** 10.1101/2025.07.27.667068

**Authors:** Alessandro D Uboldi, Sachin Khurana, Amalie A Jaywickrama, Sai Prasanna Lekkala-Lethakula, Amber Simonpietri, Karan Singh, Vinzenz Hofferek, Ushma Ruparel, Lachlan L Whitehead, Kelly L Rogers, Alexandra L. Garnham, Nichollas Scott, Nicholas J Katris, Malcolm J. McConville, David Komander, Simon A Cobbold, Christopher J Tonkin

## Abstract

*Toxoplasma* and other Apicomplexan parasites, switch between different developmental stages to persist in and transmit between hosts. *Toxoplasma* can alternate between systemic tachyzoites and encysted bradyzoite forms found in the CNS and muscle tissues. How parasites sense these tissue types and trigger differentiation remains largely unknown. We show that *Toxoplasma* differentiation is induced under glucose-limiting conditions and using a CRISPR screen identify parasite genes required for growth under these conditions. From ∼25 identified genes important for differentiation we show that lactate and glutamine metabolism is linked to differentiation and demonstrate the importance of an E3 ubiquitin ligase complex, orthologous to glucose induced degradation deficient (GID) complex in yeast and CTLH complex in humans. We show that TgGID likely regulates translational repression of a key transcription factor required for differentiation, BFD1, through its 3’ utr. Overall, this work provides important new insight into how these divergent parasites sense different host cell niches and trigger stage conversion through a ubiquitination-dependent program.

## Main Text

The phylum Apicomplexa comprises a group of obligate intracellular protozoan parasites, which include pathogens of both animals and humans. An almost universal feature of apicomplexan species is their complex lifecycles which often involve more than one host. The molecular processes which underpin key developmental steps to transition through these lifecycles, within and between hosts, at precisely the right time, are largely unknown.

A critical developmental transition in apicomplexan species is differentiation into latent forms that are able persist in their host long-term and which also resist therapeutic intervention. *Plasmodium vivax* hypnozoites can establish latent infections in the liver which are refractory to most available therapies^1,2^. *Toxoplasma gondii*, the causative agent of toxoplasmosis, establishes latent infection in humans that can persist for life. *Toxoplasma* is one of the most common foodborne parasitic diseases with over 10 million cases/yr in the USA alone, accounting for 1.2 million disability adjusted life years (DALYs)^3^. Acute toxoplasmosis is caused by tachyzoite stages, which replicate within and lyse almost any nucleated host cell causing flu-like symptoms. When tachyzoites reach muscle or CNS tissue, a proportion differentiate into latent encysted bradyzoite forms. Bradyzoites are not cleared by the immune system and are intrinsically resistant to front-line drugs, leaving a quarter of the world’s population chronically infected^4–6^. Reactivation of this latent *Toxoplasma* reservoir in immunocompromised patients (bone marrow and organ transplant recipients, AIDS patients) can be life-threatening. Similarly, latent *Toxoplasma* infections of the eye can cause retinal scarring and progressive blindness in otherwise healthy individuals with a particularly high prevalence in South America ^7,8^.

Bradyzoite differentiation is associated with transcriptional changes in hundreds to thousands of genes, which contribute to alterations in growth, morphology and metabolism of these forms ^4,9–12^. Bradyzoites divide asynchronously and grow more slowly than tachyzoites ^13,14^, express a distinct repertoire of surface antigens ^15,16^ and secrete heavily glycosylated proteins that form a fibrous cyst wall and matrix that is deposited at the parasitophorous vacuole membrane (PVM) ^11,17,18^. Bradyzoite differentiation is also associated with significant changes in central carbon metabolism. In particular, differentiation is associated with stage-specific expression of paralogous glycolytic enzymes ^19–21^ and the accumulation of amylopectin granules ^22,23^. A recent study provided evidence that bradyzoites are dependent on glycolysis and lactate fermentation for ATP generation ^24^.

Many of the changes that underlie bradyzoite differentiation are triggered by a Myb-like transcription factor, called Bradyzoite Formation Deficient 1 (BFD1)^12^. BFD1 is both necessary and sufficient for differentiation and appears to be regulated post transcriptionally, as *bfd1* mRNA is equally abundant in tachyzoite and bradyzoite stages, but BFD1 protein is not detected until differentiation has been stimulated ^12^. Subsequently, the RNA binding protein, named BFD2/ROCY (regulator of cystogenesis)^25,26^ was shown to contribute to a positive feedback loop to enhance the level of BFD1 protein expression ^25,26^.

Despite these recent insights, how *Toxoplasma* senses changes in the host environment to trigger differentiation in muscle and CNS tissue remains poorly understood. Similarly, the molecular signals that trigger BFD1 activation and parasite differentiation have not been defined. Multiple lines of evidence suggest that differentiation is triggered by host-intrinsic factors ^4,27^ as *Toxoplasma* infection of neurons and mature muscle cells stimulates spontaneous differentiation ^24,28^ ^29^. The features of these host cells that trigger *Toxoplasma* differentiation are not well understood, but differentiation has been previously tied to host cell cycle withdrawal ^30,31^, nutrient starvation and proinflammatory immune responses ^29, 32^.

### A whole genome CRISPR screen identifies genes required for utilization of glucose and glutamine

We wanted to determine which genes are required for utilization of either glucose (Glc) or glutamine (Gln) in order to understand how parasites utilize different carbon sources. We used a *Toxoplasma* CRISPR library, targeting the predicted ∼8000 genes across the parasite genome with 10 guides per gene (ie. ∼80,000 guides), and Cas9 expressing RH strain, as previously described (Fig 1A)^33^. The guide library was transfected into *Toxoplasma* and selected for a stable population in complete media (Glc^+^/Gln^+^) over 3 passages (Fig 1A). This population was then split into media that had either glucose (Glc^+^/Gln^-^) or glutamine (Glc^-^/Gln^+^) as a sole carbon source and grown for 2 subsequent passages. Guides from each population were deep-sequenced and their frequency compared between conditions (Fig 1A).

**Figure 1:**
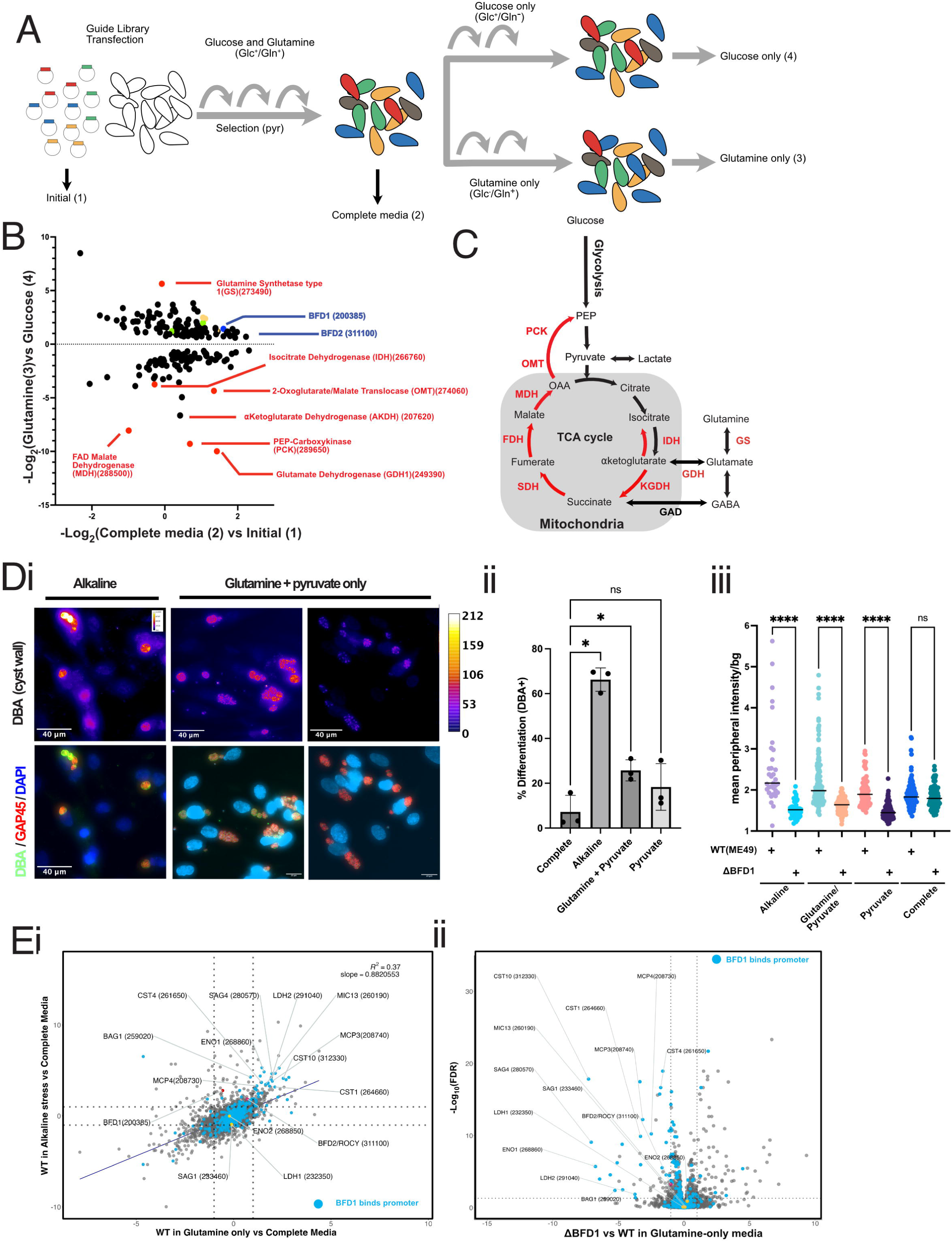
A whole genome CRISPR screen reveals that a change in carbon source trigger *Toxoplasma* differentiation. (A) Schematic representation of the experimental setup. A guide library covering the full 8000 *Toxoplasma* genes (10 guides per gene) is transfected into tachyzoites and selected for stable integration using pyrimethamine (pyr) over three lytic cycles in HFF cells in complete media (Glc^+^/Gln^+^). Populations were then transferred into media containing glucose (Glc^+^/Gln^-^) or glutamine (Glc^-^/Gln^+^) as their major carbon source and grown for a subsequent two cycles. At each stage guide pools were sampled by deep sequencing as labelled with numbers. (B) Differentially abundant hits from A (Log_2_FC (glutamine-only(sample 4) vs Glucose-only (sample 3))), graphed relative to their fitness score (Log_2_FC (Initial (sample 1) vs complete media ( sample 2) as defined by ^12^) during standard growth conditions. Genes in red highlight those predicted to be involved in central carbon metabolism (C) Schematic of *Toxoplasma* central carbon metabolism with genes identified as fitness conferring in glutamine-only conditions of CRISPR screen highlighted in red. (Di) Immunofluorescence assay of WT (ME49) and ΔBFD1 parasites infecting HFF host cells and stained with GAP45 antibodies (red)(parasite periphery) and Dolichos Bifluorus Agglutinin (DBA) (Fire tone) across Alkaline stress and when carbon source was limited to glutamine and pyruvate (Glc^-^/Gln^+^/Pyr^+^). (ii) Quantitation of *Toxoplasma* differentiation using DBA staining as a marker, comparing when parasites were exposed to complete media (Glc^+^/Gln^+^), Alkaline stress and glutamine+ pyruvate conditions (Glc^-^/Gln^+^/Pyr^+^). Mean +/− S.D plotted, n=3 biological repeats, One Way ANOVA, where **p< 0.01, ns = not significant. (iii) Quantitation of cyst wall intensity in WT and DBFD1 across conditions described in Bii. DBA intensity was calculated by measuring the mean pixel intensity of a 0.1μm thickness around the periphery of the vacuole, as marked by GAP45 and dividing this by the local mean pixel intensity of the background. Each data point represents a single parasite vacuole with the bar representing median value. Data set is a single representative experiment. Statistical testing done by ordinary one-way ANOVA where ****<0.0001, ns= not significant. (E) Quantitative Proteomic analysis of WT(ME49) and ΔBFD1 grown under conditions as marked. (i) Log-Log plot comparing changes in protein abundance when WT parasites were switched to either Alkaline Stress or Glutamine-only (Glc^-^/Gln^-^) conditions from Complete Media (Glc^-^/Gln^+^) for 48hrs. R^2^ and slope values calculated using linear regression. (iii) Volcano plot showing changes in protein abundance of WT (ME49) parental strain versus DBFD1 under glutamine as a sole carbon source. For both ii and iii; proteins are highlighted in aqua if their encoding genes promoter binds to BFD1 (as shown by ^12^) whilst proteins known to be tachyzoite-specific highlighted in yellow. Proteins known to be bradyzoite- and tachyzoite-specific are labelled.

The analysis showed that there was a good correlation between our CRISPR screen and that reported by Sidik et al, underscoring that our data was of sufficient quality and largely comparable (Fig E1A)(Table S1) ^33^. We then analyzed differentially abundant guides between Glc^+^/Gln^-^ versus Glc^-^/Gln^+^ media (Fig 1B) (Table S1). As expected, guides targeting genes involved in utilization of glucose or glutamine were differentially recovered under the two growth conditions and compared to their fitness score in complete media (Fig 1B). Interestingly, most genes that are fitness conferring in Glc^-^/Gln^+^ are largely not fitness conferring in complete media (Glc^+^/Gln^+^) (Fig 1B). Strikingly, when identified genes were mapped back onto a scheme for *Toxoplasma* central carbon metabolism, many were involved in glutamine utilization and/or predicted to be required for gluconeogenesis through to the production of PEP (Fig 1C – red). Glutamate dehydrogenase 1 (GDH1(249390)) was the highest fitness conferring gene when glutamine is supplied as the sole carbon source, followed by PEP carboxykinase (PEPCK(289650)), a predicted malate dehydrogenase (MDH(288500)) and a predicted α-ketoglutarate dehydrogenase (αKDH(207620))(Fig 1B).

To validate our findings, we made a series of transgenic *Toxoplasma* strains to both epitope tag and knock out/down genes of interest. GDH1(429390), isocitrate dehydrogenase (ICD; 266760), malate dehydrogenase (MDH;288500), Oxaloacetate-Malate Transporter (OMT; 274060) and PEP carboxykinase (PEPCK;289650) were tagged with either HA and/or the mini Auxin Inducible Degron (mAID) (Fig E1,B,C). GDH1-HA displayed a punctate and diffuse cytoplasmic localisation, whilst PEPCK-mAID-HA localized to a single puncta per tachyzoite and was associated with the mitochondrion (Fig E1B,C). *Toxoplasma* mutant lines lacking GDH1 (ΔGDH1) and conditionally regulated PEPCK (PEPCK-mAID) were made. Both grew normally as tachyzoites when cultivated in host cells in the presence of complete medium (Glc^+^/Gln^+^), but were unable to progress past the 1-2 parasite per vacuole stage when glutamine was supplied as the sole carbon source (Glc^-^/Gln^+^), suggesting that these enzymes are required to utilize glutamine as a carbon source (Fig E1C). The importance of GDH1 for glutamine catabolism was confirmed by labelling wild type and ΔGDH1-infected host cells with ^13^C- glutamine. Loss of GDH1 led to mildly increased labelling of intracellular pools of glutamine and glutamate but resulted in reduced levels of ^13^C-labeling of α-ketoglutarate (αKG) and other TCA cycle intermediates. Reduction in labeling was highly significant for αKG, malate and isocitrate and decreased (but did not reach statistical significance) for succinate, malate and citrate (Fig E1Civ). Together these data strongly implicate several enzymes in the utilization of glutamine as a carbon source during *Toxoplasma* growth in vitro.

### Loss of glucose stimulates differentiation into latent forms

Surprisingly, guides targeting BFD1 were significantly enriched when parasites were grown in Glc^-^/Gln^+^ medium, suggesting that parasites that lacked *bfd1* grew faster in this condition (Fig 1B, E1D). We hypothesized that the glutamine-only conditions (or lack of glucose) was stimulating differentiation into slower growing latent forms, and that genetic disruption of *bfd1* prevents differentiation thereby keeping them as faster growing tachyzoites, thus outcompeting other parasites in the population. Interestingly, muscle and neural tissue are thought to utilize glutamate and lactate as major carbon sources, suggesting that the Glc^-^/Gln^+^ growth conditions may mimic carbon source availability in vivo ^34^. The RH strain is known to be poor at differentiating into bradyzoite forms, thus we tested whether the cystogenic type II ME49 strain, and matched ME49 ΔBFD1 control, differentiates in glutamine and pyruvate as carbon sources (Gln^+^/Pyr^+^ medium)(addition of pyruvate was required to keep host HFF alive in these early experiments) for 48hrs. Cultivation of *Toxoplasma* in Glc^-^/Gln^+^/Pyr^+^ medium was sufficient to induce bradyzoite differentiation (DBA^+^), albeit not to the same level as control alkaline stress, as measured as a percentage of population (Fig 1Di,ii) or when the mean peripheral DBA intensity of individual vacuoles was quantitated as compared to background (Fig 1Diii). In all cases, differentiation was dependent on expression of BFD1 (Fig 1Diii).

To understand whether changes in carbon source triggered a similar genetic program to alkaline stress we quantitated changes to the proteome under these different conditions (Table S2). Growth in Glc^-^/Gln^+^ stimulated the expression of a range of proteins in a similar manner to alkaline stress (Fig 1Ei, E1E) including proteins previously shown to be induced upon bradyzoite conversion (labelled, e.g., ENO1, CST1, CST10, LDH2, BAG1) and many whose gene promoters have previously been mapped to bind to the BFD1 transcription factor and induced in a BFD1-dependent manner (Fig 1Ei-blue)^12^. Importantly, expression of all detected bradyzoite proteins in Glc^-^/Gln^+^ was dependent on expression BFD1 (Fig 1Eii).

### Sub-library CRISPR screens in vitro and in vivo identify genes required for *Toxoplasma* differentiation

These findings raised the possibility that other genes identified in our carbon-switch CRISPR screen may also be required for *Toxoplasma* differentiation into latent bradyzoite forms. To test this possibility, we filtered the CRISPR screen hits to identify guides that had the same differential abundance profile across the experiment as BFD1, identifying 60 genes. To determine which of these genes were involved in *Toxoplasma* differentiation, we synthesized a CRISPR sub-library consisting of 5 guides targeting each of these 60 genes, plus non-cutting control guides and guides that target highly fitness conferring genes (Fig 2A)(Table S1) ^33^. We first used this library to determine which genes were required for differentiation *in vitro* under standard alkaline stress conditions by transfecting the CRISPR sub-library into an ME49:Cas9 reporter line that constitutively expresses RFP protein (*gra1*_promoter_-RFP) and bradyzoite-specific mNeonGreen (mNG) protein (*bag1*_promoter_-mNG) ^12^. Parasites were then selected for stable integration of guide plasmids and then differentiated for 48 hr under alkaline stress conditions (Fig 2A). Parasite populations were then FACS sorted for mNG^+^ (bradyzoite) or mNG^-^ (tachyzoite) parasites and each sample deep sequenced to determine relative abundance of guides targeting each gene. We then compared guide frequencies in mNG^+^ and mNG^-^ back to the parental population (sample 2) to determine which genes were associated with *Toxoplasma* differentiation into latent forms *in vitro* (Fig 2B). Across our experiments, both the high fitness conferring and the non-cutting (negative control) guides performed as expected (Fig E2A). Similarly, BFD1 (dark blue) was depleted in the mNG^+^ (sample 3) vs starting population (sample 2) as compared to mNG^-^ (sample 4) population (deviation from the 1:1 line), further validating our screen (Fig 2B). A range of other genes also showed a depletion in mNG^+^ vs mNG^-^, including BFD2/ROCY (light blue), which has subsequently been shown to be important for differentiation ^25,26^(Fig 2B). Other genes with a similar depletion in mNG^+^ included those predicted to be involved in central carbon metabolism (yellow)(e.g. FNT(209800), GDH2 (293180)), RNA binding (grey)(e.g. 6xCCCH-like (301400), TTF-like (269420)) as well as ubiquitination (green)(eg OTUD1(260510), E2-H-like (300040))(Fig 2B).

**Figure 2:**
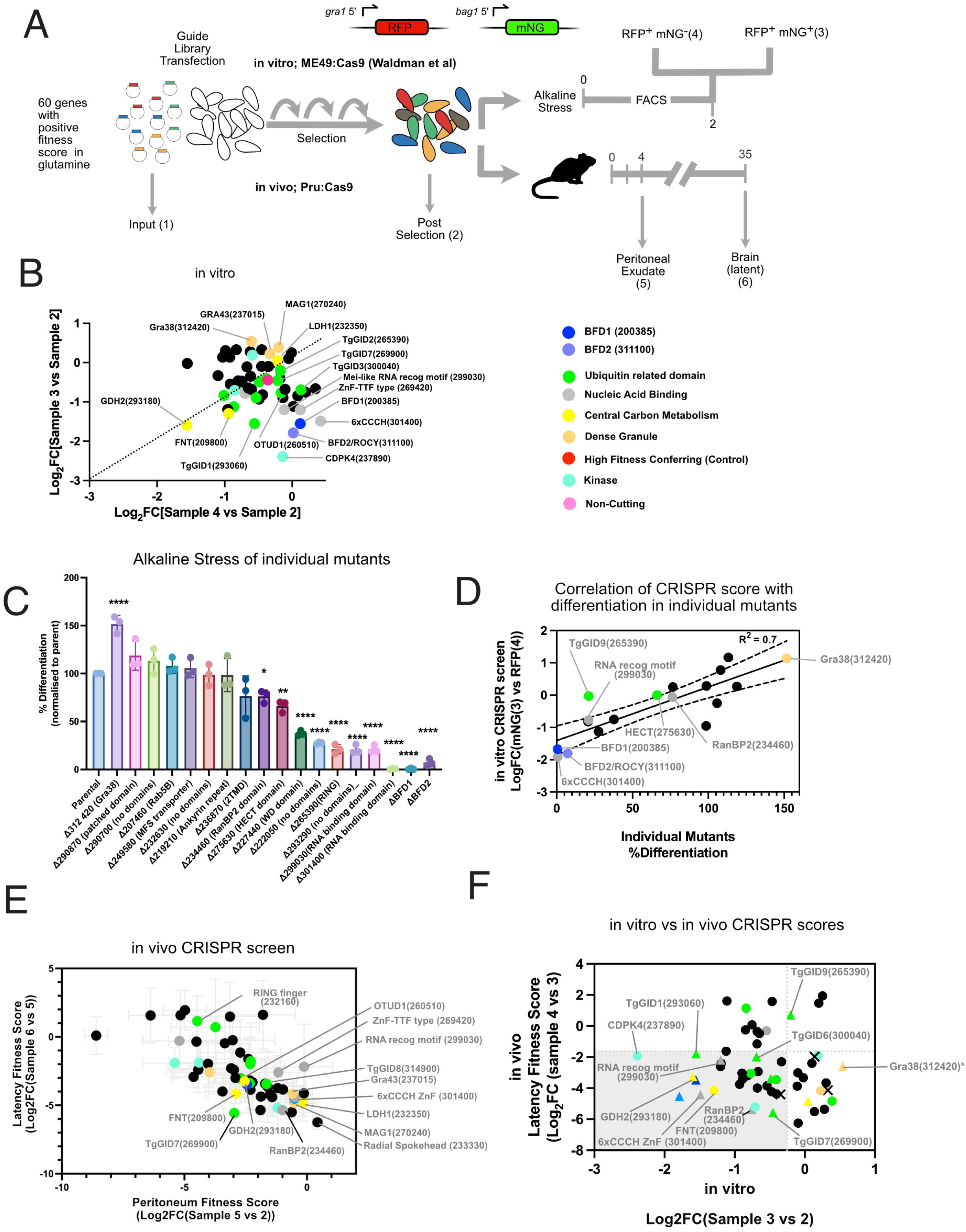
In vitro and in vivo CRISPR screens highlights genes required for differentiation of *Toxoplasma* into latent forms. (A) Schematic representation of sub-library CRISPR screens both in vitro and in vivo (mouse). Guide libraries targeting 66 genes (including controls) of 5 guides per gene, were constructed. For in vitro screen; the guide library was transfected into ME49:Cas9 reporter line, which expresses RFP constitutively using the Gra1 promoter (*gra1*_promoter_-RFP)and mNeonGreen (mNG) under control of the bradyzoite-specific BAG1 promoter (*bag1*_promoter_-mNG) ^12^, selected under pyrimethamine (pyr) for 3 cycles in HFF cells in standard conditions. Cultures were exposed to alkaline stress to induce differentiation for 48hrs and parasites were then FACS sorted for mNG expression and prepared for sequencing. For in vivo screen; the same sub-library was transfected into Pru:Cas9 and selected over 3 cycles on pyr for stable integration of plasmids and then injected into cohorts of Swiss mice. At day 4 post infection peritoneal exudate containing *Toxoplasma* was collected. At day 35 brains of mice were collected. At each step DNA was extracted and guide sequences amplified for sequencing. (B) In vitro CRISPR screen analysis; Guide frequencies at the gene level, comparing FACS sorted RFP^+^mNG^+^ (Sample 3)(y-axis) and RFP^+^mNG^-^ (Sample 4)(x-axis) back to the initial population (Sample 2). Each data point represents a single gene and are coloured based on predicted (or known) function, with corresponding key. (C) Generation of individual *Toxoplasma* mutant strains; Guide plasmids were randomly picked from sub-library and used to generate individual mutant transgenic strains in the ME49:Cas9 reporter strain^12^. Each mutant was subjected to alkaline stress for 48hrs and FACS used to analyze differentiation into *bag1*_promoter_-mNG expressing parasites. Samples were then normalized to parental strain. BFD1 and BFD2 were used as controls. (D) Correlation of CRISPR screen results with differentiation of individual mutants. R^2^ value calculated by linear regression. (E) Results of the mouse in vivo CRISPR screen, comparing Peritoneum Fitness Score (Log_2_FC (Sample 5 vs Sample 2) (x) vs Latency Fitness Score (Log_2_FC(Sample 6 vs 5))(y) Each data point represents a single gene, colour-coded as in B. (F) Comparison of in vitro and in vivo CRISPR screens, comparing the latency fitness score (as outlined in E) vs Log_2_FC(Sample 3 vs 2) (as in B). Each data point represents a single gene colour-coded as in B.

To validate further our CRISPR screen we made a series of individual mutants, by randomly picking bacterial colonies from the sub-library, sequencing and transfecting these individual guide-bearing plasmids into the ME49:Cas9 reporter line and disruption confirmed (Fig E2B)^12^. Using FACS to monitor *bag1_promoter_*-mNG we quantified differentiation (Fig 2C) we found that ΔGRA38 (312420), as compared to parental control, caused higher levels of differentiation, whilst there was no change for seven other mutants (Fig 2C). Eight mutants had a reduction in differentiation levels to varying degrees (Fig 2C). We then plotted the in vitro CRISPR scores (RFP^+^/mNG^+^ vs RFP^+^/mNG^-^) against the percentage differentiation of individual mutants and found there to be a good correlation (R^2^=0.7), providing further validation on the overall robustness of our screen and identification of genes required for differentiation (Fig 2D).

We then adapted our CRISPR screen to assess which genes were important for differentiation into latent forms in a mouse model of latent *Toxoplasma* infection (Fig 2A). We generated a Pru:Cas9 expressing strain (less virulent in mice that ME49), which we then transfected with the guide library and selected for a stable population of parasites (Fig 2A). Libraries of parasites were then injected into cohorts of Swiss mice and samples taken from the peritoneum at day 4 to assess each genes contribution to acute *in vivo* fitness. Samples were also taken from the brain at 5 weeks post-infection, a time where a latent infection has typically established (Fig 2A). Parasite samples were then deep sequenced and comparative guide frequencies measured across the experiment. A ‘Peritoneum Fitness Score’ (Log_2_FC(sample 3 vs 2)) for each gene was calculated representing the relative capacity of mutants to survive in the peritoneum (Fig E2C) as well as a ‘Latency Fitness Score’ (Log_2_FC(sample 4 vs 3) representing each genes contribution for parasites capacity to move from the peritoneum to the brain and establish a latent infection (Fig E2D). These were then combined to correlate the scores (Fig 2E). Further, we derived a scatter plot to compare results from both *in vitro* and *in vivo* screens and integrated the data on the individual mutants that we generated (triangles = defect in differentiation, X = no detected defect in differentiation) to identify the genes most likely required for differentiation (Fig 2F – grey region). Overall, this identified ∼25 new genes (plus BFD1 and BFD2 controls) as being important for bradyzoite differentiation in vitro and in vivo.

### Genes involved in central carbon metabolism are required for efficient differentiation of *Toxoplasma* into latent forms

Given our initial findings that changes in carbon source can trigger differentiation, three genes stood out to us as worthy of further investigation-– lactate dehydrogenase 1 (LDH1;232350) which interconverts pyruvate and lactate, a formate nitrite transporter 1 (FNT1;209800) which has been characterized as transporting lactate across the parasite plasma membrane ^35^ and a second predicted glutamate dehydrogenase (GDH2;293180). In each of the three genes a HA tag was added at the C-terminus to track each protein and a subsequent knockout made, each of which was confirmed by western blot (Fig E3A). We used a recently described KD3 muscle cell line, which causes spontaneous conversion into latent forms ^24^. We first measured steady-state levels of key central carbon metabolites in KD3 myotubes to see how these differed to HFF cells. Strikingly, KD3 myotubes had less steady state hexose phosphates and more glutamine and glutamic acid than HFF, mimicking conditions in the Glc^-^/Gln^+^ medium from earlier experiments (Fig 3A). We then added strains to KD3 myotubes and then fixed samples at day 2 and 5 post infection and stained for parasite cyst wall using DBA (Fig 3B). As previously described, the parental WT (ME49) strain spontaneously differentiates into bradyzoites by Day 2 with almost every vacuole having converted into encysted forms by Day 5 (Fig 3B)^24^. Whilst, both ΔLDH1 and ΔFNT1 appeared to differentiate to the same extent as WT parasites, as shown by DBA staining, some differences were observed (Fig 3B, 3C). Both ΔFNT1 and ΔLDH1 appear to have aberrant cyst walls at Day 2 (Fig 3B, grey arrowheads) and by Day 5 have an increase in smaller, DBA negative vacuoles that cluster together, typically seen when large vacuoles of parasites egress and invade neighboring cells (Fig 3B – yellow arrowheads). We also observed that cysts of ΔLDH1 parasites were also much larger than WT parasites. This data is consistent with the phenotype of these genes in the initial CRISPR screen i.e. a growth advantage that mutations provide under conditions that stimulate differentiation. Interestingly, these phenotypes were not obvious under alkaline stress conditions (Fig E3B).

**Figure 3:**
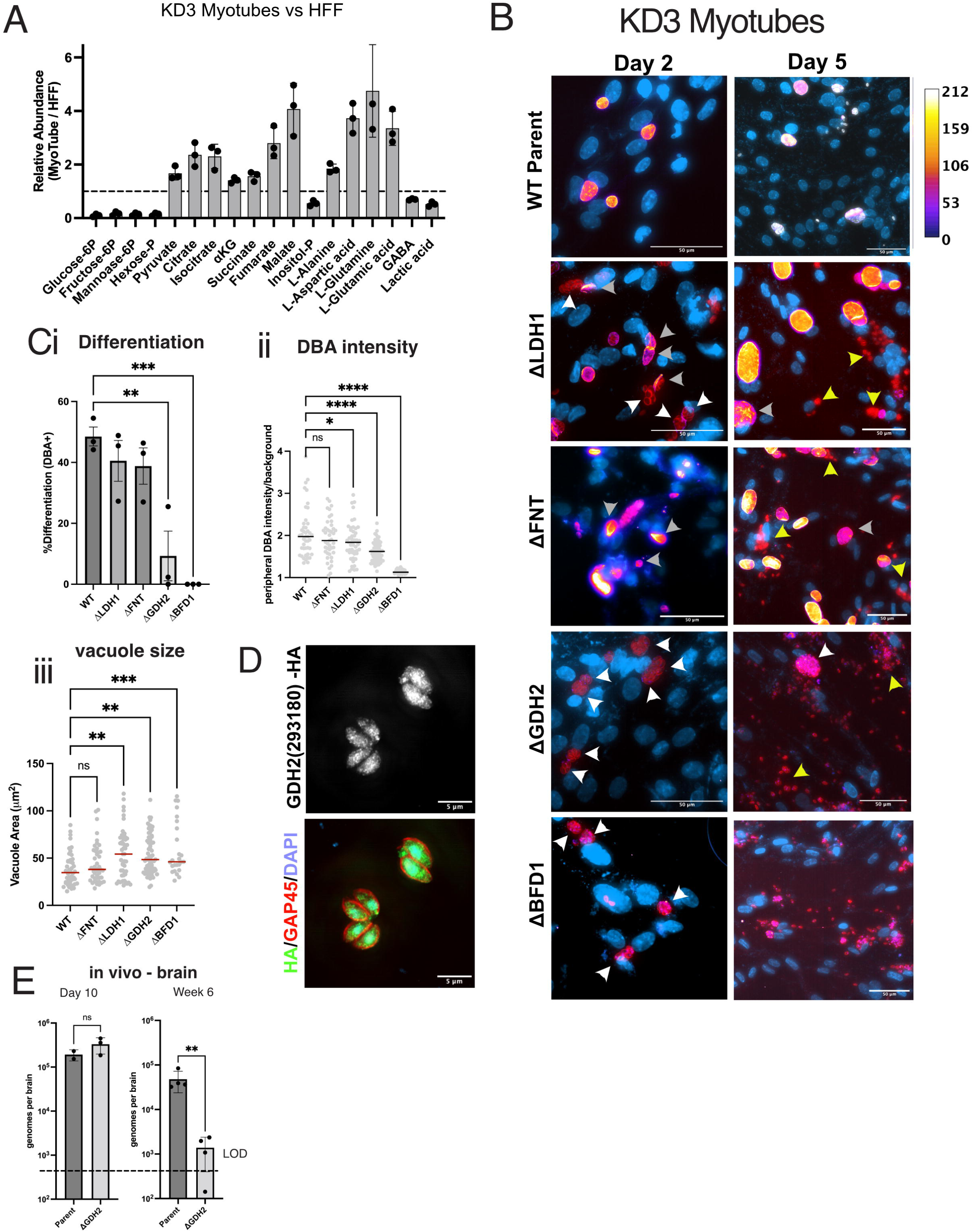
Disruption of *Toxoplasma* genes involved in central carbon metabolism causes defects in differentiation into latent forms. (A) Analysis of steady-state levels of central carbon metabolites of KD3 myotubes vs HFF, as represented as a ratio between samples, n=3 biological replicates, mean +/−SD represented. (B) Immunofluorescence of WT, ΔLDH1, ΔFNT1, ΔGDH2 and ΔBFD1 *Toxoplasma* strains infecting KD3 myotubes at day 2 and day 5 post infection. DBA(fire tones) and GAP45 (red), DAPI(blue). White arrow heads point to parasite vacuoles not stained by DBA and grey arrowheads point to disrupted morphology of cyst wall as marked by DBA. Yellow arrowheads point to small vacuoles DBA^-^ vacuoles (Ci) Percentage differentiation of ΔLDH1, ΔFNT1 and ΔGDH2 strains when grown in KD3 cells for 48hrs, as marked by DBA^+^ staining. n=3 biological repeats, mean +/−SD. Statistical testing done by ordinary one-way ANOVA where **p<0.01,***p<0.001, ns= not significant. (ii) Measurement of the cyst wall intensity across each mutant as measured by mean peripheral intensity of the DBA stain (0.1μm outside the GAP45 stained area), corrected to the mean local background. Each data point represents a single vacuole and bars represent median value. Data is a representative single biological replicate. Statistical testing done by ordinary one-way ANOVA where *p<0.05, ****p<0.0001, ns= not significant. (iii) Vacuole area in mm^2^ of mutant strains as determined by area of GAP45 staining. Data is a representative single biological replicate. Statistical testing done by ordinary one-way ANOVA where **p<0.01,***p<0.001, ns= not significant. (D) Localisation GDH2(293180)-HA by immunofluorescence in intracellular tachzyoites. (E) Analysis of ΔGDH2 vs WT parental line in mice in the brain at day 10 and week 6. Parasite numbers calculated by qPCR. n=3-4 mice, LOD = Limit of Detection. Statistical testing performed using unpaired two-tailed t-test, **p<0.01, ****p<0.0001, ns= not significant.

Quantitative analysis of differentiation of ΔGDH2 parasites showed a severe block in the formation of a cyst wall (Fig 3B,Ci,ii) and a larger vacuole area at Day 2 (Fig 3Ciii), consistent with the faster growth of this line being due to a block in differentiation. By Day 5, large ΔGDH2 vacuoles were greatly reduced in number and instead smaller clusters of DBA negative vacuoles were present, which were completely devoid of DBA staining, similar to that observed for ΔBFD1(Fig 3B). We wondered how GDH2 controls differentiation and how its function could differ from GDH1 (Fig 1). To understand this, we localized GDH2-HA by IFA and found that this enzyme is found predominantly in the nucleus (Fig 3D), suggesting that this enzyme may not function to create aKG for use in the mitochondrion, but instead could fuel nucleometabolic processes, similar to that recently described in mammalian cells ^36^. Interestingly, we could see no difference in ^13^C-glutamine incorporation in ΔGDH2 as compared to WT (Fig E3C), perhaps due to controlling a smaller pool of aKG in the nucleus as compared to the mitochondrion. Overall, these data suggest that mutations in LDH1, FNT1 and GDH2 cause parasites to grow faster in the KD3 myotube model, because of a partial or complete block in differentiation into latent forms.

We then tested whether GDH2 was required for establishment of a latent infection *in vivo*. Parental WT and ΔGDH2 lines were intraperitoneally injected into Swiss mice and infection of the brain monitored by qPCR (day 10) as well as persist into a latent infection (week 6). At day 10, when tachyzoites are typically arriving in the brain, we observed that parental WT and ΔGDH2 strains were in equal amounts (Fig 3E). However, by week 6, when most parasites are present as encysted bradyzoites, much fewer parasites were detected for ΔGDH2 as compared to the WT (Fig 3E). Together this work suggests that perturbations in enzymes which are predicted to be important for the synthesis or metabolism of glutamine and lactate have impacts on *Toxoplasma* differentiation into latent forms and thus links these two processes.

### *Toxoplasma* contains a GID E3 ubiquitin ligase complex

We identified ubiquitin machinery^37^, including several components of the GID/CTLH-type E3 ubiquitin ligases as important for differentiation in both screens^38,39^. In yeast, GID is responsible for the degradation of gluconeogenic enzymes when glucose is present ^40,41^. In Drosophila, the CTLH E3 complex (and Marie Kondo E2) is required for the maternal zygotic transition (MZT) early in development, where it degrades proteins to relieve translational repression of RNAs ^42,43^. In higher eukaryotes, the CTLH complex controls erythropoiesis, organ development and metabolic enzyme levels ^44^ ^45,46^. We genetically tagged TgGID1,7,8 and 9 with HA, and localized these within intracellular tachyzoite stages and confirmed tagging by western blot (Fig 4A,E4A). showing a punctate pattern through the parasite’s cytosol, and some higher density of staining in the nucleus (Fig 4A, E4Ai). To determine whether *Toxoplasma* indeed has a GID complex, we performed immunoprecipitation (IP) experiments with each of these lines (Fig 4B, EB). Pull-down of each component resulted in co-precipitation of the other 4 members, strongly suggesting that *Toxoplasma* does indeed have a GID complex (Fig 4Bii)(Table S2). In addition, we identified another complex component – TgYPEL5 – which has been shown to be a negative regulator of substrate binding ^46^. Genetic tagging and IP of TgYPEL5-HA confirmed that this is a bone fide member of the complex (Fig E4B). This suggests that *Toxoplasma* has a GID complex. We found no evidence of GID4 and 5, which acts as substrate receptors in yeast^47^, however, *Toxoplasma* does contain a GID7 orthologue, which can also act as a substrate receptor and is responsible for formation of a GID-chelate for recognition and ubiquitination of homo-tetramers (Fig4Biii) ^48^.

**Figure 4:**
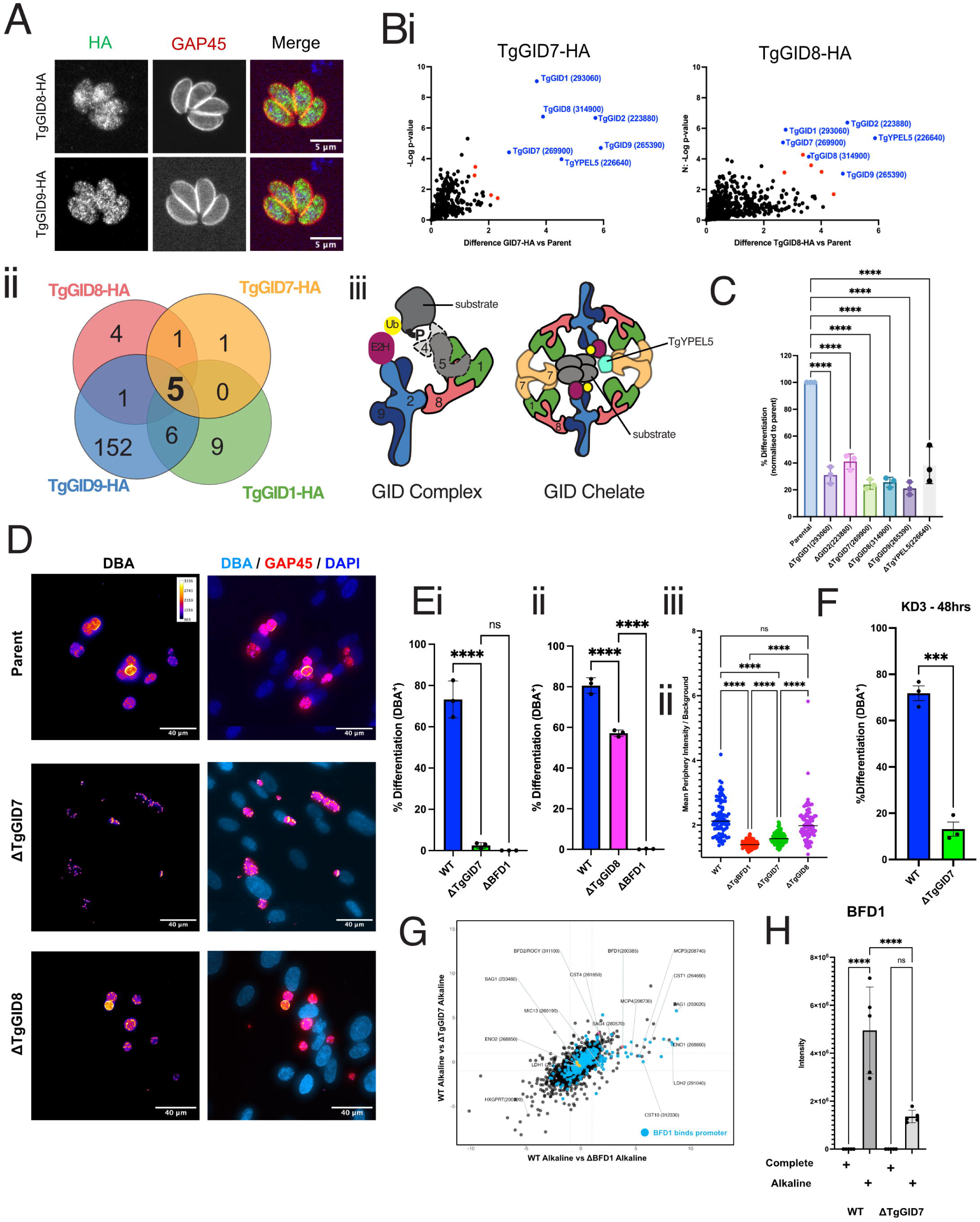
*Toxoplasma* contains a GID E3 ubiquitin ligase complex which is required for differentiation into latent forms. (A) Localisation of TgGID7-HA and TgGID8-HA by immunofluorescence together with GAP45 (parasite periphery) and DAPI (Bi) Quantitative αHA Immunoprecipitation of TgGID7-HA and TgGID8-HA as compared with parental control. Statistically significant hit marked in red and other TgGID components in blue. (ii) Venn diagram comparing results across all four experiments highlighting the identification of 5 common proteins. (iii) a model of TgGID complex based on previous structural data ^46,48,72^. (C) Analysis of differentiation in individual CRISPR mutant lines generated in ME49:Cas9 reporter strain ^12^ as determined by BAG1_promoter_-mNG by FACS. Normalized to parental strain. n=3 biological replicates, One way ANOVA, where ****p<0.0001. (D) Representative images of IFA analysis of differentiation after 48hrs of alkaline stress of parent, DTgGID7 and DTgGID8 strains, where DBA(Fire tone) marks cysts wall, GAP45 marks parasite periphery and DAPI-host cell nuclei. (E) Analysis of differentiation of mutants after 48hrs of alkaline stress (i) Percentage differentiation of WT (parent) versus ΔTgGID7 and ΔBFD1 as determined by DBA cyst wall staining. n=3 biological replicates, mean +/− SD, One way ANOVA, where ****p<0.0001, ns=not significant. (ii) Percentage differentiation of WT (parent) versus ΔTgGID8 and ΔBFD1 as determined by DBA cyst wall staining. n=3 biological replicates, mean +/− SD, One way ANOVA, where ****p<0.0001. (iii) Cyst wall intensity measurements across WT, ΔBFD1, ΔTgGID7 and ΔTgGID8 as determined by mean peripheral DBA staining intensity (0.1μm outside GAP45 staining) divided by the local mean background intensity of individual parasites. Single representative experiment, where each datapoint represents a single vacuole and bar represents median value. One way ANOVA, where ****p<0.0001 and ns=not significant. (F) Analysis of differentiation of parent and ΔTgGID7 in KD3 myotubes as quantitated by DBA staining. n=3 biological replicates, mean+/− SD, Unpaired, two-tailed t-test, where ***p<0.001. (G) Log-Log plot of quantitative global proteomic analysis of WT vs ΔBFD1 vs ΔTgGID7 upon 48hrs of alkaline stress. Blue datapoint represent proteins, where their gene promoter has been shown to bind to BFD1 ^12^. Yellow datapoints represent known tachyzoite-specific proteins. Annotated proteins represent known bradyzoite and tachyzoite-specific proteins. Dotted lines represent Log_2_FC=1in both dimensions. (H) BFD1 protein levels as determined by quantitative proteomics in WT parent and ΔTgGID7 strains in both complete media (standard tachyzoite growth conditions) and alkaline stress. n=5 biological repeats, mean +/−SD. One way ANOVA, where ****p<0.0001 and ns=not significant.

### TgGID is required for differentiation of *Toxoplasma* into latent forms

To determine if TgGID is required for differentiation into latent forms we generated mutant populations lacking the six identified genes using ME49:Cas9 reporter strain (Fig 4C, Fig E4C) and then assayed their capacity to differentiate into latent forms using alkaline stress for 48hrs. Differentiation was quantitated by FACS (*bag1_promote_*_r_-mNG) as compared to WT (Fig E4Ciii). TgGID mutants could only differentiate at 20-40% of the level of the WT parental line, as measured by mean fluorescent intensity of mNG (Fig 4C). We then made clonal knockout lines of TgGID7 and TgGID8 (Fig E4D) in a WT ME49 strain and quantitated differentiation by imaging after staining the cyst wall with DBA (Fig 4D,E,F). Both ΔTgGID7 and ΔTgGID8 were defective in differentiation under both alkaline stress conditions (Fig 4D,E) and when glutamine was the sole carbon source (Fig E4E). We then measured vacuole area (μm^2^) as a surrogate for growth rate under both alkaline and glutamine-only differentiation conditions. Whilst there was no statistical difference in area in alkaline stress between either mutant compared with WT and ΔBFD1, there was under glutamine-only conditions (Fig 4Eiii,iv) suggesting a growth advantage, thus helping to explain their initial identification in the genome-wide CRISPR screen (Fig 1). ΔTgGID7 was also defective in differentiation in the KD3 muscle cell model (Fig 4F, E4F).

To understand what role TgGID plays during *Toxoplasma* differentiation into latent forms we performed quantitative proteomics on ΔTgGID7 as compared to the WT parent and ΔBFD1 in complete (Glc^+^/Gln^+^) medium and under alkaline stress for 48hrs (Table S2). Interrogation of all TgGID components upon knockout of TgGID7 showed lower relative levels as compared to WT parent, suggesting that deletion of this component destabilizes the whole complex (Fig E4Gi). Principal component analysis (PCA) shows tight clustering of biological replicates and similarity between ΔTgGID7 as compared to ΔBFD1 in alkaline stress conditions (Fig E4Gii). To further understand similarities and differences between loss of ΔTgGID7 and ΔBFD1 we compared both these parasite strains to the parental WT under alkaline stress conditions (Fig 4G). We found that loss of ΔTgGID7 (vs WT) resulted in a similar loss of expression of known and predicted bradyzoite proteins, as compared to loss of BFD1, albeit at slightly reduced levels (Fig 4G). This is further exemplified by comparing intensity measurements of SAG1 and CST1, canonical markers of tachyzoites and bradyzoites, respectively, across all conditions (Fig E4Giii, iv). Interestingly, ΔTgGID7 parasites also appeared to have a lower the amount of BFD1 protein under 48hrs of alkaline stress, suggesting a link between this E3 ubiquitin ligase and regulation of this transcription factor (Fig 4H).

### TgGID can regulate differentiation through the 3’ utr of *bfd1*

To characterize further a potential link between the BFD1 and the TgGID complex we generated knockouts of both TgGID7 and TgGID8 in the previously described inducible BFD1 strain – DD-BFD1/DBFD1 (Fig 5A)^12^. This DD-BFD1 strain was constructed by genetic deletion of endogenous *bfd1* gene using a *sag1* promoter driven mNeonGreen (*sag1_prom_*-mNG) as a selectable marker, followed by integration, at the *hxgprt* locus, of another copy of BFD1 fused to the destabilization domain (DD) (DD-BFD1), driven by the tubulin promoter (Fig 5Bi). In this strain, the addition of the synthetic compound Shield-1(Shld-1) stabilizes DD, preventing its proteosomal turnover, allowing for the fused BFD1 protein to accumulate and differentiation to be induced (Fig 5Bi)^12,49,50^. Upon genetic removal of either TgGID7 or TgGID8 and addition of Shld-1 we found that both mutations led to defective differentiation, as measured by DBA staining (Fig 5A). Interestingly, during this analysis we noticed that every knockout clone of TgGID8 expressed high levels of mNG, whilst the parental (and sister clones that were not knockouts) (DD-BFD1 WT) were only weakly fluorescent (Fig 5Bii,C). Given that mNG is controlled by the *sag1* promoter region we wondered whether this transgene was acting as a surrogate for changes in SAG1 protein levels, a canonical tachyzoite antigen. We showed this was true by correlating mNG levels with anti-SAG1 staining by IFA across a population of DD-BFD1/ΔTgGID8-Shld parasites (Fig E5A). We also observed that the DD-BFD1 parental line, in the absence of Shld-1, had small puncta of CST1 staining (Fig 5Bii - white arrows), typical of parasites at early stages of differentiation, which were mostly lost upon deletion of TgGID8 (Fig 5Bii). We therefore hypothesized that suppression of the mNG signal and punctate CST1 staining in DD-BFD1 strain was due to some ‘leaky’ BFD1 protein being present in the absence of Shld-1. Furthermore, we hypothesized that loss of TgGID8 suppresses this leaky level of DD-BFD1, thus allowing for upregulation of *sag1_promoter_*-mNG and turning off expression of small amounts of CST1.

**Figure 5:**
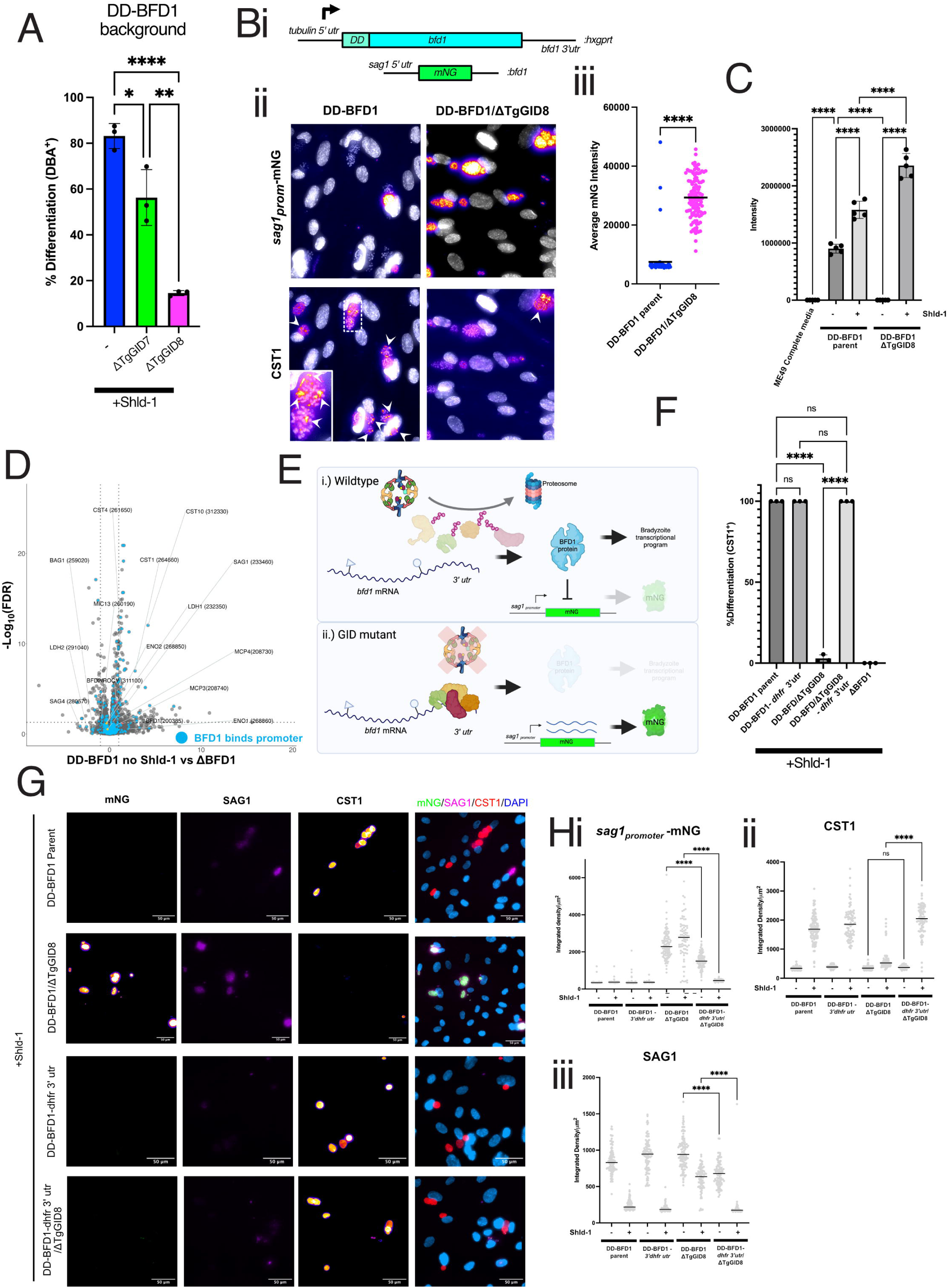
TgGID likely acts through the 3’ utr of *bfd1* to regulate differentiation. (A) Quantitation of differentiation into latent forms of DD-BFD1 parent versus ΔTgGID7 and ΔTgGID8 after 48hrs treatment with Shield-1 (Shld-1) compound. n=3 biological repeats, mean +/−SD, One way ANOVA, where *p<0.05, **p<0.01 and ***p<0.0001 (Bi) Schematic representation of DD-BFD1 strain genotype as previously constructed ^12^. *sag1_promoter_*-mNG was used to genetically disrupt the endogenous *bfd1* gene and an ectopic copy of bfd1 is fused to the destabilization domain (DD), controlled by the *tubulin_promoter_* and the *bfd1* 3’ utr. (ii) Representative IFA images of DD-BFD1 parent versus DD-BFD1/ΔTgGID8 (without Shld-1), showing levels of mNG and CST staining. Inset and white arrowheads demonstrate punctate CST1 staining. CST1 and mNG (fire tone) and DAPI (greyscale).(iii) Quantitation of mNG signal from DD-BFD1 parent and DD-BFD1/ΔTgGID8 (without Shld-1 treatment) as measured by mean pixel intensity of individual vacuoles by IFA. Single representative experiment, where each datapoint represents a single vacuole and bars represent median value. Unpaired, two tailed t-test, where ****p<0.0001. (D) BFD1 protein levels as determined by quantitative global proteomics. n=5 biological replicates, mean +/− SD, One way ANOVA, where ****p<0.0001. (E) Volcano plot comparing global protein abundance as measured by quantitative proteomic analysis. Protein-encoded genes whose promoter binds to BFD1 are highlighted in red. Known tachyzoite-specific proteins in yellow. Proteins that are known to be specific to bradyzoites or tachyzoites annotated. Horizontal dotted line represents FDR=0.05 and vertical dotted lines represent Log_2_FC= 1. (F) Speculative model of TgGID function through the 3’utr of bfd1 mRNA transcript. (H) Quantitative IFA analysis of percentage differentiation as marked by CST1 antibodies of *bfd1* 3’utr swap lines as compared to parental and ΔBFD1. n=3 biological repeats, mean+/− SD, One way ANOVA, where ****p<0.0001, ns=not significant (G) Quantitation of mNG, SAG1 and CST1 staining of *bfd1* 3’ utr swap strains and their parental controls (i) Representative images showing staining of mNG, SAG1 and CST1 staining patterns. Quantitation of pixel intensity values for mNG(ii) SAG1(iii) and CST1 (iv) across all samples. In each case data shown is from single representative experiment, where each datapoint represents a single vacuole and bars represent median value. One way ANOVA, showing select comparisons, where ****p<0.0001 and ns=not significant.

To test this, we first performed global quantitative proteomics on DD-BFD1 parent, DD- BFD1/ΔTgGID8 and ΔBFD1 strains both with and without (+/−) Shld-1(Table S2). Like deletion of TgGID7, loss of TgGID8 in the DD-BFD1 background also showed lower relative levels of other TgGID components (Fig E5Bi). PCA showed good clustering of biological replicates and separation of conditions (Fig E5Bii). Importantly, addition of Shld-1 in DD-BFD1 strain lacking TgGID8 impedes upregulation of known and predicted bradyzoite proteins (Fig E5Biii). As hypothesised, we could show that, even in the absence of Shld-1, this transcription factor can still be detected, as compared to the undetectable levels in WT ME49 parasites in complete media (Fig 5C). Furthermore, comparison of DD-BFD1-Shld-1 with ΔBFD1 showed that this leaky expression leads to the upregulation of bradyzoite proteins, including CST1 (Fig 5D). As expected, addition of Shld-1 almost doubles the level of DD-BFD1 protein, and in line with our hypothesis, the loss of TgGID8 reduces the level of DD-BFD1 protein down to an undetectable level. Interestingly, loss of TgGID8 leads to a greater amount of DD-BFD1 protein, in the presence of Shld-1 (Fig 5C). The reason for this, is not understood, but appears in contrast to what happens to BFD1 upon loss of TgGID7 in ME49 WT strain (Fig 4H). We also tested whether loss of TgGID8 had an impact on the levels of *bfd1* mRNA by qPCR but found little to any effect, suggesting that changes in BFD1 protein abundance, after the loss of TgGID, was likely occurring largely post-transcriptionally (Fig E5C).

We wondered whether TgGID could be mediating the translational repression of *bfd1* through its 3’ utr (which is present in the DD-BFD1 transgene) (Fig 5Bi)^12^, whereby this E3 ubiquitin ligase complex can ubiquitinate and degrade proteins bound to this regulatory element allowing for BFD1 translation to occur (Fig 5E). This hypothesis is based on precedence for translational repression being crucial for developmental processes in related *Plasmodium* spp which acts through the 3’ utr of target genes ^51,52^. Furthermore, Drosophila orthologous CTLH E3 ubiquitin ligase is known to relieve translational repression by ubiquitinating and degrading RNAbinding proteins ^42,43^. In this model loss of TgGID would therefore prevent the release of translational repression and lead to diminished BFD1 protein, which then cannot suppress *sag1_promoter_*-mNG nor activate downstream genes for differentiation to take place (Fig 5E). To test this, we wondered whether replacing the *bfd1* 3’ utr with the 3’ utr of the *dhfr* housekeeping gene would overcome the block in differentiation imposed by loss of TgGID. We hypothesized that this would remove the translational repressive effects of proteins that bound to *bfd1* 3’utr that were under control of TgGID proteosomal degradation. To test this, we replaced the 3’ utr of *bfd1* in DD-BFD1 in parental and ΔTgGID8 strains to the 3’ utr from the housekeeping gene *dhfr*. We then subjected parasites to treatment with Shld-1 for 48hrs and used imaging as a readout for differentiation using CST1 antibodies to mark the cyst wall. We found that swapping to the *dhfr* 3’ utr in the parental DD-BFD1 line had no measurable impact on the differentiation capacity, nor did it affect the levels of CST1, SAG1 or *sag1_promoter_*-mNG with or without Shld-1, as compared to the parental DD-BFD1 strain (Fig 5G,H,I). However, when the *bfd1* 3’ utr was swapped for the *dhfr* 3’ utr in the DD-BFD1/ΔTgGID8, differentiation was reinstated (Fig 5G,H). Furthermore, after the addition of Shld-1, swapping the 3’ utr’s led, once again, to the suppression of the *sag1_promoter_*-mNG reporter (Fig 5Ii), reinstated the upregulation of CST1 (Fig 5Iii), and reversed the defective downregulation of SAG1 (Fig 5Iiii). In the absence of Shld-1, spots of CST1 once again became visible in DD-BFD1/ΔTgGID8-*dhfr* 3’ utr similar to that observed in the DD-BFD1 parental strain (Fig E5D)(as described in Fig 5Bii). To ensure this effect was not due to large changes in *bfd1* mRNA, transcript was measured via qPCR. Whilst there were some mild changes observed under no Shld-1 conditions, there were no significant changes in +Shld-1 samples (Fig E5C). Overall, this data provides a genetic link between TgGID and the 3’ utr of *bfd1* and suggests a role for this E3 ubiquitin ligase in translation control of master transcription factor that controls *Toxoplasma* differentiation into latent forms.

## Discussion

Apicomplexan parasites differentiate through multiple cell types during their complex life cycles, allowing these parasites to be transmitted between hosts and alternate between acute and long-term latent infections. Despite their importance, the molecular processes underpinning how Apicomplexan parasites sense their environment and coordinate differentiation remains poorly understood. Here we have exploited CRISPR screens to identify genes required for *Toxoplasma* differentiation in vitro and in vivo. We provide evidence that *Toxoplasma* bradyzoite differentiation can be triggered by changes in carbon source availability in the host cell and is genetically linked to parasite central carbon metabolism. We also provide evidence for a role for ubiquitination in *Toxoplasma* differentiation through a GID/CTLH-type E3 ubiquitin ligase complex and show that this likely regulates BFD1 protein levels of this key transcription factor.

Changes in tissue type and host environment have a profound effect on all pathogens, allowing them to undergo genetic programs of adaptation, trigger a developmental switch to progress to the next life cycle stage, or set up a latent infection for long term persistence in the host. In this study we provide both genetic and biochemical evidence that *Toxoplasma* can use changes in carbon source availability to trigger bradyzoite differentiation. We show that cultivation of infected host cells in glucose-limited/glutamine replete media triggers differentiation in a similar manner to alkaline stress, the most widely used method for activating differentiation in *Toxoplasma*. Whilst alkaline stress is unlikely to be a physiological trigger, changes in carbon catabolism are known to differ in cell types where *Toxoplasma* spontaneously differentiates into latent forms, such as muscle and neurons. These cell types are thought to be less reliant on glucose to fuel their central carbon metabolism, but rather can use lactate and glutamate, both of which occur at high levels in these tissues ^34^. Indeed, we show that KD3 human muscle cells, that cause the spontaneous differentiation of *Toxoplasma* into bradyzoites, have lower steady state levels of hexose phosphates and higher levels of glutamate and glutamine than HFF, a cell type which allows very low levels of spontaneous differentiation, consistent with this hypothesis.

Further supporting of a link between central carbon metabolism and *Toxoplasma* differentiation into latent forms is our identification of three genes predicted to be required for utilization of exogenous lactate and glutamine – lactate dehydrogenase 1 (LDH1), formate nitrate transporter 1(FNT1)(a lactate transporter) and a second copy of glutamate dehydrogenase (GDH2). Individual mutants of these three genes all appear to have defects in differentiation to varying degrees when grown in the human KD3 myotubes. ΔLDH1 and ΔFNT1 mutants formed abnormal cyst walls, grew faster, and egressed from their host cell more readily that WT, potentially signifying more frequent de-differentiation into tachyzoites or defects in complete or stable formation of slower-growing latent bradyzoite forms. This suggests that these enzymes, through either the production, transport or utilization of lactate are required for the correct establishment of latent forms. This work is in line with previous observations that LDH1 plays a role in differentiation under alkaline stress and is defective in formation of latent stages in the brain ^53,54^.

We also show that GDH2 partially localizes to the parasite nucleus and mutants of this enzyme were completely incapable of differentiation into latent forms, suggesting that the interconversion of glutamine to a-ketoglutarate (αKG) is important for differentiation of *Toxoplasma*. Interestingly, it has recently been reported that a GDH in mammalian cells creates αKG in the nucleus, which is then used as a cofactor for both DNA and histone demethylase enzymes ^36^. This provides a tantalizing hypothesis as to how central carbon metabolism in *Toxoplasma* could be linked with differentiation. Indeed, there is little understood about the role of epigenetic regulation in the development of latent bradyzoite stages, despite their being a precedence for this in sexual stage development in both *Toxoplasma* and related *Plasmodium falciparum* parasites ^55–58^. Similarly, epigenetic regulation required for *P. falciparum* sexual development is linked to lysophosphatidylcholine metabolism, providing further evidence for a metabolic link to epigenetic regulation in this group of parasites ^59,60^.

Our CRISPR screens revealed approximately 25 genes were highly likely to be required differentiation in vitro and in vivo, including genes predicted to bind RNA and/or DNA, two kinases, a series of hypothetical proteins with unknown function and several genes that encode proteins with ubiquitin-related domains. Five of these genes encode proteins that form part of GID/CTLH-type E3 ubiquitin ligase complex and are important for differentiation. Furthermore, the screen also identified an E2 enzyme orthologous to E2H in yeast and Marie Kondo in Drosophila which donates charged ubiquitin to the GID and CTLH E3 ligases, respectively. Yeast GID, Drosophila and Human CTLH’s have all been shown to form K48 chains and target proteins for degradation and thus it is highly likely that this is also true for TgGID. Consistent with this notion, an OTU-like deubiquitinase (DUB), was also identified in the screen, which we have recently shown can degrade K48 chains ^37^.

What is not clear at this stage, is how substrates are recognized by TgGID as we could not identify an orthologue to either GID4 or GID5 in our immunoprecipitation experiments under either standard or differentiation inducing conditions or with homology based-searches. In humans, recognition of some substrates does not require GID4, instead, GID7/WDR26 can bind substrates through an internal KRKW motif ^46^. This is interesting as GID7/WDR26 is also responsible for assembling the GID chelator structure which allows for recognition of a subset of substrates. Biochemical analysis also suggests that the GID7/WDR26 binding interface can be blocked by YPEL5, which contains a N-terminal motif that mimics the substrate binding motif ^46^. Given that TgGID appears to have both a GID7 and YPEL5 orthologues, but apparently lacks GID4 and 5, this suggests that this mode of substrate recognition is most likely.

We provide genetic evidence that TgGID potentially regulates BFD1 translation through the 3’ utr. In other systems, including related-*Plasmodium* parasites, the 3’ utr of specific RNAs can bind protein complexes that repress their translation ^51,52^. Furthermore, in Drosophila the orthologous CTLH complex ubiquitinates and degrades translational repressive machinery bound to key regulatory RNA, allowing for translation to occur ^42,43^. Our data is consistent with the 3’ utr of *bfd1* mRNA also binding translationally repressive machinery and that TgGID acts to degrade these proteins to promote the synthesis of BFD1 protein. What these proteins are, however, is not yet known, but it is possible that this machinery could include other proteins that we have identified in our CRISPR screen, including one or more of the RNA binding proteins.

Our work led to the subsequent identification and characterization of a GID complex in related malaria-causing *P. falciparum* parasites, which is presented in an accompanying manuscript (Marapana et al). Here, the authors describe the important role PfGID plays in differentiation of *P. falciparum* into its sexual stages. Furthermore, Marapana et al highlight that PfGID likely controls the release of translational repression by acting to degrade a key RNA binding protein. Together, both studies suggest that similar mechanisms relying on ubiquitination as a central regulatory process to control differentiation between life cycle stages across all Apicomplexa, perhaps at multiple different developmental points, of this important group of intracellular eukaryotic parasites. The identification of GID/CTLH complex in Apicomplexa and its role in cell fating adds to the growing level of evidence as to the importance of ubiquitination during developmental processes and fate-decisions including yeast, humans, Drosophila and now apicomplexans.

## RESOURCE AVAILABILITY

### Lead contact

Chris Tonkin

### Materials availability

Plasmids and transgenic *Toxoplasma* parasite lines generated in this study are available from the lead contact upon request.

### Data Availability Statement

Proteomics datasets are available through the following codes: Project accession: PXD058492, Token: YMpN3bJ6vH6h, Project accession: PXD058495, Token: ySmymGTvo2rt, Project accession: PXD058547, Token: 7hi1QAUgl3Am. The Macro developed for analysing the imaging data is available at https://github.com/BioimageAnalysisCoreWEHI/tonkin_lab/blob/main/cTonkin_radial_inten sity_measure.ijm.

## ACKNOWLEDGEMENTS

We are deeply indebted to the laboratory of Sebastian Lourido (Whitehead Institute, MIT, USA) for sharing of reagents both for the CRISPR screen as well as several parasites strains critical for this study. Thanks also go to Dr Laura Dagley for management of the proteomics facility at WEHI and Dr Stephen Wilcox for help with sequencing. This work was funded through grants from the National Health and Medical Research Council of Australia (NHMRC) (GNT2029730) and The National Institute of Health USA (R01AI172823). This work was also supported by the Victorian State Government Operational Infrastructure Support grant (Institutional grant) and Australian Government NHMRC IRIISS.

## AUTHOR CONTRIBUTIONS

CJT and ADU conceptualized the studies, wrote the draft, and all authors contributed to data analysis as well as reviewing, editing and approving the manuscript. ADU and UR performed CRISPR screens. ALG analysed the resulting data. ADU, SK AAJ constructed transgenic parasites and together with SPLL and AS performed phenotypic analysis. SK, KS, NS and SAC performed and analysed proteomic datasets. LKW and KLR wrote code, provided access to equipment to performing quantitative imaging. NJK, VH and MJM performed and analysed metabolic tracing experiments. DK provided key expertise and reagents for proteomic experiments.

## Materials and Methods

### Experimental Model and Subject details

#### Parasite and host cell culture

*Toxoplasma gondii* parasites were propagated in confluent monolayers of Human Foreskin Fibroblasts (HFFs). HFFs were grown to confluency in host cell medium (DMEM (Gibco) supplemented with 2 mM Glutamax (Gibco), 1 X penicillin/streptomycin (Gibco) and either 10% heat inactivated Cosmic Calf Serum (Hyclone) or 10% FBS (Sigma). For infection with *Toxoplasma*, this medium was removed from HFF monolayers and replaced with the same medium, but containing 1% FBS, referred to as ‘complete medium.’

For differentiation of *Toxoplasma* in HFFs, tachyzoites were allowed to infect cells typically with a multiplicity of infection of 1 parasite per 10 host cells. The infected cells were returned to culture for 2 hours, after which the medium was replaced with either alkaline differentiation medium (RPMI (Gibco)-20 mM HEPES, pH 8.1; 5% FBS; 25 mM glucose; 2 mM Glutamax) or glutamine differentiation medium (Glucose-free DMEM (Gibco), 6mM Glutamax; 2 mM glutamine; 2 mM sodium pyruvate; 1% dialyzed FBS) and returned to culture at 37 °C + 10% CO_2_ for glutamine differentiation, or CO_2_-free conditions for alkaline differentiation. Differentiation medium was replaced with fresh medium daily for the duration of the differentiation period.

#### KD3 Human Muscle cells

The culturing and maintenance of the KD3 immortalized human muscle cell line and infection with *Toxoplasma* protocol was adapted from previous work ^61^. The progenitor myoblast cells were grown in a humidified incubator at 37°C and 10% CO_2_ and cultured in DMEM containing 4 mM L-glutamine, 1 mM sodium pyruvate, 1X penicillin/streptomycin, supplemented with 25 mM D-glucose, 2% Ultroser G (Sartorius) and 20% heat-inactivated fetal bovine serum (FBS). The myoblasts are maintained sub confluent during passage in T75 flasks and are induced into myotubes in 6-well or 24-well plates on coverslips at 70%-80% confluency. Myogenesis is induced by the media being changed into DMEM containing 4 mM L-glutamine, 1 mM sodium pyruvate, 1X penicillin/streptomycin, supplemented with 25 mM D-glucose, 2% donor horse serum, 10 µg/mL human recombinant insulin, 5 µg/mL human holo-transferrin, and 10 nM sodium selenite forming myotubes ready for infection on days 5-7 post induction.

*Toxoplasma* differentiation media referred to as ‘cyst media’ is made up of RPMI 1640 7.4 pH containing 4 mM L-glutamine and supplemented with 5 mM D-glucose, 50 mM HEPES, 2% donor horse serum, 10 µg/mL human recombinant insulin, 5 µg/mL human holo-transferrin, 10 nM sodium selenite, and 1X penicillin/streptomycin. Plates were washed with room temperature PBS and are replaced with cyst media prior to infection. *Toxoplasma* lines were maintained on HFF monolayers in between experiments and a dilution of 7.2 x 10^3^ tachyzoites per cm^2^ were prepared in cyst media and added to each well. Plates were moved directly into an ambient CO_2_ incubator at 37°C and the non-invaded parasites washed off after 24 hours and replaced with fresh cyst media. For extended differentiation the media was be replaced every 2 to 3 days leaving 25% volume of the prior media to avoid complete disruption of the monolayer.

#### Mice

Mice were bred and maintained at The Walter and Eliza Hall Institute of Medical Research Animal Facility. All experiments were approved by the institute Animal Ethics committee and performed in accordance with the Australian Code for the Care and Use of Animals for Scientific Purposes and the National Health and Medical Research Council of Australia Code of Practice for the Care and Use of Animals for Scientific Purposes guidelines. Mice were housed in individually vented cages in a specific pathogen free facility with a 12 hr light/dark cycle and unlimited access to chow and water. 6–8-week-old female mice were infected by intraperitoneal injection with the indicated number of tachyzoites. Mice were humanely euthanized using CO_2_ exposure at required time points.

### Method Details

#### Primers and plasmids

Oligonucleotide primers were from IDT and cloning was carried out by Q5 mutagenesis (NEB), HiFi DNA assembly (NEB), or In-Fusion cloning (Takara). Details of primers and plasmids used in this study can be found in Table S4.

#### Parasite transfection

For standard transfections approximately 15 μg of guide plasmids and 15 μg of homology repair template PCR products were combined and ethanol precipitated, before resuspending in 20 μl of Lonza P3 solution + Supplement solution. *Toxoplasma* infected HFFs were scraped with a cell scraper (Corning) and passed through a 27-gauge needle to release tachyzoites. The tachyzoites were spun down at 11,000 x *g* for 4 min, and the medium removed. Parasites were resuspended in the DNA mixture and electroporated using a Lonza nucleofector, using program F1-115.

#### Generation of Transgenic strains

##### Preparation of Cas9-expressing RH:Dhxgprt parasites and Pru:Dhxgprt parasites

To create the RH:*cas9* strain, RH:Δ*hxgprt* parasites ^62^ were transfected simultaneously with pCas9-CAT (Addgene # 80323) and pU6-Decoy (Addgene #80324) plasmids together, as done previously ^33^. To create Pru:*cas9*, Pru:Δ*hxgprt* parasites were transfected with linearized pBM019 (Addgene #128179) plasmid that encodes Cas9 ^63^. In both cases, stable integration of the plasmids was achieved using selection with 20 mM chloramphenicol (Sigma). Expression of Cas9 was assessed by IFA using anti-FLAG M2 antibody (Sigma).

##### Generation of guide plasmids for gene disruption and epitope-tagging

To generate guide plasmids for gene disruption and epitope-tagging with either 3xHA or 3xFLAG-mAID tags, Q5 mutagenesis (NEB) with pSAG1-Cas9-GFP-U6-sgUPRT (Addgene #54467) DNA template was used ^64^. The oligonucleotides used to create guide plasmids for epitope tagging and knocking out various genes of this study are shown in S4 (oligonucleotides 25 - 80). Successful mutagenesis PCR was assessed by agarose gel electrophoresis, before treating 1 μl of mutagenesis PCR products with KLD reaction mix (NEB). The KLD reactions were transformed into DH5alpha bacteria and guide plasmids produced were sequenced by Sanger sequencing to confirm the presence of the expected guides within the plasmids.

Homology repair templates were prepared by PCR using oligonucleotides comprised of 40bp of homology to the gene of interest and 20bp of homology to a region in one of several plasmids (Table S4), available on request. The oligonucleotides used to create homology repair template DNAs are listed in Table S4 (oligonucleotides # 81- 116). Prime Star Max DNA polymerase was used to amplify the homology repair templates.

##### Creation of parasite strains in ME49:Dku80: Dbfd1:DD-BFD1-Ty background

To create Δ*gid8* in parental ME49:D*ku80*:D*bfd*1:DD-BFD1-Ty:*bfd1* 3’UTR parasites (previously created by Waldman et al, 2020(^12^), pSAG1-Cas9-U6-sg*gid8* guide plasmid and homology repair template with phleomycin selection cassette were used for transfection. To replace the BFD1 3’UTR in ME49:Δ*ku80*:DD-BFD1-Ty line with a *dhfr* 3’utr, two guide plasmids (created by Q5 mutagenesis using oligonucleotides 77-80; Table S4), one targeting each end of the *bfd1* 3’UTR, and a repair template encoding the *dhfr* 3’UTR (created using oligonucleotides 115 and 116; Table S4) were combined and used for transfection. The plasmid pLIC-3HA-HX (a kind gift from Vern Carruthers) was used as the template for the DHFR 3’utr. Since the homology repair *dhfr* 3’utr PCR product did not contain a drug selection cassette, two days after transfection, the parasites were cloned out and clones with 3’utr’s successfully swapped from *bfd1* 3’utr to *dhfr* 3’utr were identified by genotyping using oligonucleotides #x and y (Table S4).

#### Whole genome CRISPR screen

Three biological replicates of a whole genome CRISPR screen were carried out as described previously by Sidik et al., 2016 ^33^ with some modifications. For each replicate, eight 60 ug aliquots of whole genome CRISPR library plasmid DNA (Addgene #80636) were linearized by AseI (NEB) digestion, ethanol precipitated, air dried and resuspended in 131 ml of cytomix (10 mM KPO4; 20 mM KCl, 5 mM MgCl_2_, 25 mM HEPES, 2 mM EGTA, pH 7.6 with KOH). Following precipitation ∼ 50 mg DNA was recovered for each aliquot. To each DNA-containing tube, 8 ml of 100 mM ATP (Sigma) and 8 ml of 250 mM glutathione (Sigma) was added. RH:Δ*hxgprt*:*cas9* parasites were prepared from eight T150 flasks (Corning) of infected HFFs. The parasites obtained were combined in complete medium, passed through 27-gauge needles (Becton Dickinson) and centrifuged at low speed (35 x *g*) for 5 min to pellet intact cells and large debris. Parasites in the supernatant were aliquoted into eight Eppendorf tubes at 8.7 x 10^7^ parasites/tube. Each tube housing parasites was washed once with 500 ml cytomix, before resuspending parasites in 245 ml of cytomix. The DNA/ATP/glutathione mixture was added to the parasites, followed by 8 ml of 7.5 mM CaCl_2_. Eight transfections were carried out in 4 mm gap cuvettes (BTX) using the BTX square wave electroporator (Havard Apparatus) with the following settings: 1.7 kV; two 176 ms pulses; 100 ms intervals. Immediately following each transfection, the contents of the cuvettes were made up to 40 ml with D1 medium, and 5ml was added to each of eight fresh T150 flasks containing 30 ml D1 medium/40 mM chloramphenicol (to maintain Cas9 expression) and cultured overnight.

The next day, the medium was removed from the flasks and the infected monolayers were washed 3 times with 8 ml PBS, before adding 30 ml fresh complete medium containing 1 mM pyrimethamine (Sigma) and ∼10U of Turbo DNAse (Ambion) and returned to culture. After ∼ 24 hours of culture, the cells/parasites were scraped with a cell scraper (Corning), needle passed through 27-gauge needles and prepared as described above, before adding 5.75 x 10^7^ parasites to each of eight new T150 HFF flasks containing 30 ml D1 medium plus 1 mM pyrimethamine to select for parasites with stably integrated guide plasmids. Two additional passages with pyrimethamine selection were carried out, following which the stable parasite population was split into two populations of 6.9 x 10^7^ parasites. These were washed once with either glutamine-only medium (glucose-free DMEM (Gibco), 6mM Glutamax; 2 mM glutamine; 1% dialyzed FBS) or glucose-only medium (glutamine-free DMEM (Gibco), 25 mM glucose; 1% dialyzed FBS), before adding the parasites to six T150 HFF flasks, each containing 30 ml of the same media. The following day the media were removed from the flasks, the infected monolayers washed 3 times with PBS and replaced with 30 ml fresh media. After ∼ 24 hours, the resulting parasites were added to six new T150 HFF flasks with glutamine-only (Glc^-^/Gln^+^) and glucose-only (Glc^+^/Gln^-^) medium and returned to culture. A second cycle of growth in glutamine-only Glc^-^/Gln^+^) and glucose-only (Glc^+^/Gln^-^) medium and was carried out, as above. The final parasite yields for parasites grown in glucose-only medium and glutamine/GlutaMax-only medium were aliquoted into Eppendorf tubes and stored at −80°C until required for genomic DNA preparation and sequencing.

##### Guide sequencing

For each biological replicate, genomic DNA was prepared from the pyrimethamine-selected population, glucose-selected population and glutamine/GlutaMax-selected parasite population using phenol-chlorofom-isoamyl alcohol (Sigma). PCR reactions were then performed to amplify the guide region from the genomic DNAs as well as from the original CRISPR library (Addgene #80636) and amplified library used for transfections. PrimeStar Max DNA polymerase (Takara) and unique combinations of barcoded primers 1 – 20 (Table S4) were used to amplify the guide region. For PCR amplification of the library, 60 ng of DNA was used as template, while for the CRISPR screen genomic DNA populations, two PCR reactions were performed and combined, each with 300 ng of genomic DNA as template. PCR cycling conditions used to amplify the library were: 98 °C for 30 s, 30 cycles of 98 °C for 10 s, 63 °C for 5 s and 72 °C for 5 s, followed by 72 °C for 2 min. For amplification of genomic DNA, the following conditions were used: 98 °C for 30 s, 35 cycles of 98 °C for 10 s, 55 °C for 5 s and 72 °C for 1 min) followed by 72 °C for 5 min. The resulting PCR products were purified and 5 ml aliquots from each sample were pooled, and a fraction of the pooled DNA was purified using a Bioline PCR purification kit and subjected to next generation sequencing using a MiSeq kit (Illumina) to identify the guide sequences.

#### CRISPR screens with sub-library

##### Generation of CRISPR sub-library

Sixty genes were selected from the initial whole genome CRISPR screen as being potentially involved in *Toxoplasma* differentiation by filtering using fold-change and statistical cut-offs. In the first step, genes were selected that had no statistical difference between complete medium (Glc^+^/Gln^+^) and glucose-only medium (Glc^+^/Gln^-^) in calculated fitness scores (Log_2_FC). In the second step, genes were further filtered for those that had statistically significant positive Log_2_FC values when comparing glutamine-only (Glc^-^/Gln^+^) with glucose-only (Glc^+^/Gln^-^) conditions. The first 5 guides of each of the 60 genes, as defined by Sidik et al., 2016 ^33^, were then synthesized (CustomArray) with appropriate overhangs to allow for Gibson cloning and assembly into pU6-DHFR as described previously by Sidik et al., 2018 ^33^ with some modifications.

To create the sub-library, plasmid pU6-DHFR (Addgene #80329) was digested twice overnight at 37 °C with BsaI-V2 (NEB). The second BsaI-V2 digestion Antarctic phosphatase (NEB) was added for 2 hours at 37 °C and purified after running the digested plasmid DNA on an agarose gel. At the same time, the sub-library was PCR amplified in two steps from a larger library of pooled oligonucleotide guide sequences made by CustomArray. PrimeStar Max DNA polymerase and primers 21 and 22 (Table S4) were used to amplify the sub-library guide sequences from 20 ng of pooled library template using a gradient PCR with annealing temperatures of 55, 57, 59, 61 and 63 °C. The gradient PCR reactions were pooled and purified using a PCR purification kit (Bioline). A second gradient PCR was performed to generate ends for insertion into *Bsa*I-digested pU6-DHFR using primers 23 and 24 (Table S4) and 2 ng of purified pooled PCR product from the first reaction as template. Following PCR, the reactions were combined and run on an agarose gel. The band of around 70 bp was cut out from the gel and purified with a gel purification kit. HiFi DNA assembly master mix (NEB) was used to assemble the sub-library into the pU6-DHFR *Bsa*I-digested added to backbone using a 1:5 molar ratio of plasmid vector to PCR products. Following the assembly reaction, DH10β high efficiency competent cells were transformed with the assembly reaction to produce the sub-library, which was amplified as described by Sidik et al.,2018 ^65^.

##### In vitro screen with sub-library

Three biological replicates of the screen were performed. For each replicate, six aliquots of 30 μg of *Ase*I-digested CRISPR sub-library were electroporated into aliquots of 1 x 10^8^ ME49; *gra1 5’*-RFP, *bag1 5’*-mNG parasites (previously created by Waldman et al, 2020 ^12^) using a BioRad square wave electroporator (BioRad) with the following settings: 1.7 kV; two 180 µs pulses; 5 s intervals. Immediately following each transfection, the contents of the cuvettes were made up to 30 ml with complete medium, and 5ml was added to each of six T150 HFF flasks containing 30 ml complete medium and placed in the incubator overnight. The following day, the medium/uninvaded parasites were aspirated off, and the infected monolayers were washed once with complete medium, before adding 30 ml of complete medium containing 1 μM pyrimethamine (Sigma) and 10 U of Turbo DNase (Ambion) and returning to culture. After 9 days in culture (when most cells in the monolayer were full of vacuoles that were egressing or ready to egress), the parasites/cells were scraped and centrifuged at 872 x *g* for 15 min. The cells/parasites were resuspended complete medium and passed through 27-gauge needles twice. Parasites were then spun at 35 x *g* for 5 min to pellet large cell debris. Finally, parasites in the supernatants were pelleted by centrifugation at 872 x *g* for 5 min and added to 6 fresh T150 HFF flasks containing 30 ml complete medium and 1 μM pyrimethamine and returned to culture. After 2 days in culture, the infected monolayers from the six T150 flasks were scraped and processed as above. A third selection with pyrimethamine was carried out as above. Of the total number of parasites obtained, 5.5 x 10^7^ parasites were added to each of four T150 flasks, and the remainder were frozen at −80 °C. The 4 flasks were returned to culture for 4 hours, after which the medium was aspirated off, the monolayers washed once with alkaline differentiation medium, after which the medium was replaced with alkaline differentiation medium and placed in an incubator lacking CO_2_. The following day, the media were removed from the flasks and replaced with fresh media, before returning to culture.

After 48 hours of growth in alkaline differentiation medium, the infected cells were treated with trypsin-EDTA (Gibco), and the cell suspension was combined with 25 ml of PBS/5% FBS and centrifuged at 872 x *g* for 10 min to pellet cells/parasites. The cells/parasites were then resuspended in 10 ml PBS/5% FBS, followed by centrifugation at 872 x *g* for 5 min. The parasite/cell pellets were then resuspended in PBS/2% FBS at 10, 000 cells per ml in FACS tubes and sorted on a AriaFusion cell sorter (BD Biosciences) to yield RFP/mNeonGreen (mNG^+^) and RFP-only (mNG^-^) populations. Cells infected with RFP-expressing tachyzoites and differentiated mNG-expressing parasites were used as controls for FACS parameters. The sorted parasites/cells were transferred to low bind Eppendorf tubes and centrifuged at 12 000 x *g* for 5 min. Pelleted cells/parasites were then frozen at −80 °C. Genomic DNA was prepared for pyrimethamine-selected parasites and parasites differentiated in alkaline conditions (mNG^-^ and mNG^+^ populations). The guides for these populations as well as for the amplified CRISPR sub-library used for the screen were PCR amplified and sequenced by next generation sequencing, as described above for the whole genome CRISPR screen.

##### In vivo sub-library screen

Three biological replicates of the *in vivo* sub-library screen were performed as per Guiliano et al 2024 ^66^ with some modifications. For each replicate, eight electroporation reactions of 60 μg of *Ase*I-digested BFD1-like CRISPR library into 5 x 10^7^ parasites of Pru:*cas9* parasites were carried out. A BioRad square wave electroporator was used with the following settings: 1.7 kV; two 180 µs pulses; 5 s intervals. Immediately following transfection, the contents from all 8 cuvettes were made up to 40 ml with complete medium and 5ml was added to each of eight T150 flasks containing 30 ml complete medium + 40 µM chloramphenicol to maintain Cas9 expression and returned to culture. The following day, the medium was removed from the 8 flasks and the infected monolayers were washed with PBS, before 30 ml of complete medium containing 1 μM pyrimethamine and 10U of Turbo DNAse was added to each flask and returned to culture. After ∼ 6 days of growth, the cells/parasites from the eight T150 flasks were scraped and spun down in 50 ml tubes for 20 min at 872 x *g*. The parasites obtained from every 2 x T150s flasks were resuspended in 10 ml complete medium and needle passed through 27-gauge needles twice, before centrifuging at 35 x *g* for 5 min to pellet debris. The supernatants were centrifuged at 872 x *g* and 2 x 10^7^ of the resultant parasites were added to each of eight T150 HFF flasks containing 30 ml complete medium and 1 μM pyrimethamine, before returning to culture at 37 °C. The following day, the medium containing uninvaded/dead parasites was removed by aspiration and the monolayers washed once with PBS, before adding 30 ml of fresh complete medium containing 1 μM pyrimethamine to each flask and returning to culture at 37 °C until pyrimethamine-resistant parasites were produced. After 3 rounds of pyrimethamine selection, parasites were pelleted and washed in PBS prior to intraperitoneal injection of mice.

Six 6-to-8-week-old female C57/BL6 mice were each infected with 5,000 parasites (for acute time points) and 20 mice were infected with 600 parasites (for chronic infections). The remaining parasites from the pyrimethamine-selected population were frozen at −80 °C. After 4 days of acutely infected mice were euthanized and peritoneal exudate lavages were collected. The exudates in PBS were centrifuged at 872 x *g* for 5 min to pellet cells/parasites before resuspending the parasites/cells in 1 ml complete medium. Half of the exudate volumes were then used to infect HFFs in T25 flasks to amplify the number of parasites present within the exudates, while the other half were pelleted and frozen at −80 °C. The parasites were expanded in T25 flasks until the parasites had completely lysed the host cell monolayer, collected and frozen at −80 °C - the time to achieve this was variable for each individual mouse exudate.

After 5 weeks, brains were extracted from chronically infected mice that were euthanized. The brains were placed inside wells of 12-well plates (Corning) containing 500 μl complete medium and a 3 ml syringe plunger was used as a pestle to homogenize the brain material. The brain material was then passed through a 100 μm strainer (Corning) together with 4 ml of complete medium. The brain homogenate was then needle-passed twice through 27-gauge needles and spun down at 872 x *g* for 5 min. The pellets were added to T25 HFF flasks for expansion. Once the parasites had completely lysed the host cell monolayer, the expanded parasites were separated from host debris and brain material by passage through a needle twice and spinning at low speed (35 x *g* for 5 min) to remove debris/cells. The parasites were then pelleted by centrifugation at 872 for 5 min, washed once with PBS and stored at −80 °C.

Parasite aliquots from pyrimethamine-selected parasites, parasites obtained from exudates of mice infected acutely for 4 days and parasites obtained from brains of mice chronically infected with *Toxoplasma* for 5 weeks, as well as from the amplified CRISPR sub-library used for transfection, were utilized for genomic DNA preparation and guide sequencing. The guides present in these populations were PCR amplified and sequenced by next generation sequencing using a MiSeq 300cyc Micro one kit (Illumina).

##### CRISPR screen data analysis

Raw counts for each guide RNA (sgRNA) were collected from two data files, merged based on matching barcodes, and processed to remove low-abundance guides. Annotation for each guide was added using gene names. Quality control (QC) steps included checking the total number of guides detected per sample, the proportion of guides with zero counts, and the total counts. Filtering was applied, removing genes with a count of less than 10 in at least 3 samples. To account for differences in sample composition, counts were normalized using the TMM method^67^. Batch effects were assessed and corrected using the removeBatchEffect function from the Limma package^68,69^. Differential expression (DE) was assessed for pairwise comparisons using quasi-likelihood negative binomial generalized log-linear models fitted with the glmQLFit function from the edgeR package^67^, adjusting for replicate effects. Empirical Bayes quasi-likelihood F-tests identified genes with significant changes (up-regulation or down-regulation), and the false discovery rate (FDR) was controlled at 5%^70^. MDS plots ^68^ and heatmaps were generated to visualize sample similarity and gene expression patterns.

##### Immunoblotting

Freshly egressed tachyzoites were lysed using 150µl HEPES, 150mM NaCl, 1mM MgCl_2_, 1% v/v NP-40, supplemented with benzonase nuclease (Merck) and EDTA free protease inhibitors (Roche) for 60 minutes while kept on ice. Samples were then boiled for 5 minutes with 1X Laemmli sample buffer (containing 200mM Tris pH 6.8, 8% SDS, 40% glycerol, 0.05% bromophenol blue) with 5% b-mercaptoethanol and centrifuged at 13,000 rpm. Appropriate volumes of the sample were either resolved on 4-12% SDS-PAGE gels or 3-8% Tris Acetate gels depending on respective molecular weight of the proteins. Proteins were then transferred on a nitrocellulose membrane for 60 min at 100 V while using a western transfer buffer containing 25 mM Tris, 200 mM glycine and 20% v/v methanol. The nitrocellulose membrane was then blocked in 5 % w/v skimmed milk in 0.1 % Tween-20 followed by primary and secondary antibody exposure using 2.5% w/v skimmed milk in 0.1 % Tween-20. The blots that were probed with IRDye 800CW α-rat, IRDye 800CW α-mouse and IRDye 680RD α-rabbit antibodies were imaged using the Odyssey Clx imager (LICORbio). The blots that were probed with chemiluminescent antibodies conjugated to HRP were imaged on the ChemiDoc system (Biorad). The brightness and contrast were adjusted in FIJI software (ImageJ).

##### Immunoprecipitations and mass-spectrometry

Tachyzoites cultured in a T25 flask containing human foreskin fibroblasts (HFFs) were harvested and passed through a 27-gauge needle followed by centrifugation at 2000rpm for 5 minutes at room temperature. The parasites were then washed 3X with ice cold PBS and were either utilised immediately or frozen at −80°C. Parasites were lysed using 150µl HEPES, 150mM NaCl, 1mM MgCl2, 1% v/v NP-40, supplemented with Benzonase (Merck) and EDTA free protease inhibitors (Roche) for 60 minutes while kept on ice. The parasite lysate was then centrifuged at 13,000 rpm at 4°C for 10 min and the supernatant was then incubated with anti-HA agarose beads (Roche) overnight at 4°C. The proteins and the antibody coupled agarose beads were then washed 5X with ice cold lysis buffer containing 0.1% v/v NP-40 and the proteins were then eluted with appropriate concentrations of elution buffer containing SDS.

Immunoprecipitated proteins bound to beads were boiled in 4% SDS, 10mM Dithiothreitol, 100mM Tris pH 8.5 at 90 °C for 10 minutes with shaking at 2000 rpm and then alkylated with 40 mM iodoacetamide in the dark for 1 hour. The resulting lysate was then prepared for digestion using S-traps mini columns (Protifi, USA) according to the manufacturer’s instructions. Samples were acidified to 1.2% phosphoric acid and diluted with seven volumes of S-trap wash buffer (90% methanol, 100 mM triethylammonium bicarbonate pH 7.1) before being loaded onto S-traps and washed 3 times with S-trap wash buffer. Samples were then digested with 2μg of Trypsin (1:100 protease:protein) in triethylammonium bicarbonate pH 8.5 overnight at 37°C before being collected by centrifugation with washes, followed by 0.2% formic acid followed by 0.2% formic acid / 50% acetonitrile. Samples were dried down and further cleaned up using C18 Stage tips to ensure the removal of any particulate matter before being dried by vacuum centrifugation.

C18 enriched proteome samples were re-suspended in Buffer A* (2% acetonitrile, 0.01% trifluoroacetic acid) and separated using a two-column chromatography setup composed of a PepMap100 C18 20-mm by 75-µm trap (Thermo Fisher Scientific) and a PepMap C18 500-mm by 75-µm analytical column (Thermo Fisher Scientific) using a Dionex Ultimate 3000 UPLC (Thermo Fisher Scientific). Samples were concentrated onto the trap column at 5 µl/min for 6 min with Buffer A (0.1% formic acid, 2% DMSO) and then infused into an Orbitrap 480™ (Thermo Fisher Scientific) or a Orbitrap Q-Exactive plus at 300 nl/minute via the analytical column. Peptides were separated by altering the buffer composition from 3% Buffer B (0.1% formic acid, 77.9% acetonitrile, 2% DMSO) to 23% B over 59 minutes, then from 23% B to 40% B over 10 minutes and then from 40% B to 80% B over 5 minutes. The composition was held at 80% B for 5 minutes before being returned to 3% B for 10 minutes. The Orbitrap 480™ Mass Spectrometer was operated in a data-dependent mode automatically switching between the acquisition of a single Orbitrap MS scan (300-1600 m/z, maximal injection time of 25 ms, an Automated Gain Control (AGC) set to a maximum of 300% and a resolution of 120k) and 3 seconds of Orbitrap MS/MS HCD scans of precursors (normalise collision energy of 30%, a maximal injection time of 60 ms, a AGC of 250% and a resolution of 15k). The Orbitrap Q-Exactive Mass Spectrometer was operated in a data-dependent mode automatically switching between the acquisition of a single Orbitrap MS scan (375-1400 m/z, maximal injection time of 50 ms, an Automated Gain Control (AGC) set to a maximum of 3e6 and a resolution of 70k) and 15 Orbitrap MS/MS HCD scans of precursors (stepped normalise collision energy of 28%; 30% and 32%, a maximal injection time of 110 ms, a AGC of 2e5 and a resolution of 35k).

Identification and LFQ analysis were accomplished using MaxQuant (v1.6.3.4 or 2.4.7.0)^52^ using the T. gondii GT1 proteome from ToxoDB [3] (Version 48, 8460 Entries, downloaded 14/10/2020) with Carbamidomethyl (C) allowed as a fixed modification and Acetyl (Protein N-term) as well as Oxidation (M) allowed as variable modifications with the LFQ and “Match Between Run” options enabled. The resulting data files were processed using Perseus (version 1.6.0.7)^53^ with missing values imputed based on the total observed protein intensities with a range of 0.3 σ and a downshift of 1.8 σ and statistical analysis undertaken using a two-tailed unpaired T-test. The mass spectrometry proteomics data have been deposited to the ProteomeXchange Consortium via the PRIDE^54^ partner repository with the dataset identifier:TgGID8-HA VS WT IPs (Project accession: PXD058492, Token: YMpN3bJ6vH6h), TgGID9-HA (CTLH-HA) and TgGID1-HA (SPRY-HA) IPs (Project accession: PXD058495, Token: ySmymGTvo2rt), TgGID7-HA and TgYPEL5-HA IPs (Project accession: PXD058547, Token: 7hi1QAUgl3Am)

#### Metabolomics Analysis

Parasites were grown in regular culture medium (low glucose DMEM 1%FBS) on confluent HFF cells for 72 hours until cultures were fully lysed. Extracellular parasite cultures were filtered with a 3μm polycarbonate filter and spun down at 2000rpm for 5 minutes at room temperature. Parasite pellets were washed once with DMEM-lite (DMEM with no glucose, glutamine, pyruvate, phenol red) and spun as before. Parasite pellets were resuspended in DMEM-lite to a concentration of approximately 5.0x10^7^ cells ml^-1^(2x). The parasite suspension (0.5 ml) was added to 0.5ml of DMEM-lite containing a 2x concentration of ^13^C_5_-glutamine (2mM) with pyruvate (2mM, natural abundance), resulting in 1ml of a 1x solution (2.5 cells ml^-1^, 1mM ^13^C glutamine, 1mM pyruvate). The parasite suspension was incubated for 4x hours at 37°C, before parasites were metabolically quenched by briefly (5 sec) immersing the tube in a dry ice/ethanol bath and then transferring it to an ice bath. Parasites were pelleted by centrifugation (8000rpm, 4°C for 2 min), and pellets washed twice with cold PBS, before being stored at −80 until extraction.

HFF and myotubes were cultured under standard conditions for metabolomic analyses. The culture medium was removed by aspiration and the host cell monolayers washed with ice-cold 1xPBS before being placed on ice. Cells were suspended in PBS by scraping, and pelleted by centrifugation (2000rpm, 4°C for 5 min). Pellets were aspirated and washed twice with 1xPBS before storage in −80°C.

Cell pellets were extracted in 250μL of ice-cold MTBE:MeOH (3:1 v/v), and sonicated for 15 min, prior to the addition of 125μL of H_2_O : MeOH (3:1 v/v) containing internal standards. The mixture was shaken (1000rpm, 4°C, 30 min, Thermomixer), then centrifuged in a microfuge (15,000g, 4°C, 15 min) to pellet insoluble material and separate organic/polar phases. The bottom phase was transferred to a 96-well plate and dried down under N_2_ gas. Extracts were resuspended in either methanol:water (1:2 v/v) for Ion-Chromatograph Mass Spectrometry (ICMS)) or acetonitrile:H_2_O (4:1 v/v) for HILIC-MS. Peaks were picked using Elmaven software. Isotope normalization was done using an application built in Shiny around the package IsoCorrectoR which is available on BioConductor repository ^71^. Y-axes represent %^13^C incorporation. Graphs with an identical ME49 control were derived from datasets from the same experiment sharing the same controls.

##### Immunofluorescence assays

Infected HFF monolayers grown on cover slips in wells of 6-well plates were fixed with PBS/4% formaldehyde for 20 min. The fixative solution was removed, and parasites washed once with PBS. This was followed by permeabilization for 10 min with 0.1% Triton X-100/PBS. The wells were then washed once with PBS and blocked for at least 1 hour at room temperature with PBS/3% BSA. Primary antibodies diluted in PBS/3% BSA were then added to the wells and incubated for at least 1 hour at room temperature or overnight at 4 °C, before washing four times for 5 min in PBS. Secondary antibodies diluted in PBS/3% BSA were added to the wells for at least 1hr at RT in the dark. Samples were then washed four times for 5 minutes in PBS, with DAPI (5ug/ml) added in the penultimate wash step. Samples on coverslips were then mounted with Vectorshield and sealed with nail polish.

##### Imaging and quantification

Imaging was performed on a Zeiss Axioscope equipped with Zen software. All images were deconvolved and processed in ImageJ. In the case of quantitative imaging a custom macro was developed to allow for measuring both area and pixel intensity (https://github.com/BioimageAnalysisCoreWEHI/tonkin_lab/blob/main/cTonkin_radial_inten <u>sity_measure.ijm</u>). Briefly, the macro would allow researchers to annotate regions of interest around specific parasites which would be automatically thresholded and a periphery region defined by a user specified distance inward from the parasite boundary. Areas and intensities were measured in the “inner” region, the periphery, and the background to allow for normalisation when required. Radial distribution of intensity was also optionally recorded. Output values from the macro were then plotted, analyzed and statistics calculated in GraphPad.

##### Preparation of parasites for global proteomics

Parasites were grown in HFFs for ∼ 40 hours before being made to egress by scraping the monolayers and passing the cells/parasites through 27-gauge needles twice. A low-speed centrifugation of 35 x *g* for 5 min was used to pellet cellular debris. The parasites remaining in the supernatant were added to T25 flasks (5 replicate flasks for each strain) containing complete medium at an M.O.I of ∼1 parasite per host cell. After approximately 4 hours, uninvaded parasites were washed off and replaced with either complete medium, alkaline differentiation medium or glutamine/GlutaMax/pyruvate-medium lacking glucose. The following day the media in the flasks were replaced with fresh media and returned to culture. After 24 hours the infected HFF monolayers were scraped from the flasks and needle passed twice through 27-gauge needles. Debris were removed by centrifugation at 35 x *g* for 5 min. Supernatants were then centrifuged at 872 x *g* for 5 min to pellet parasites. Parasites were resuspended in 1 ml PBS in Protein low bind Eppendorf tubes and spun at 11,000 x *g* in a microfuge for 4 min to pellet parasites - the pellets were stored at −80 °C until required.

The pellets obtained were digested in 5% SDS in 50 mM triethylammonium bicarbonate (TEAB) pH 8.5. Lysate volume for equal protein amount was calculated from the bicinchoninic acid assay (BCA) results conducted on aliquots using Thermo Scientific™ Pierce™ BCA Protein Assay Kits. Lysates were reduced using 120 mM TCEP, alkylated using 500 mM chloroacetamide (CAA), and acidified using phosphoric acid diluted to 27.5%. Acidified lysates were then mixed with S-trap washing buffer (100 mM TEAB in 90% methanol, pH 7.55) and subsequently loaded onto the S-traps. Samples were washed 3 times with S-trap washing buffer and digested using trypsin and Lys-C in TEAB pH 8.5, with 1 μg of protease per 10 μg of protein. The proteins were digested for 1.5 hours at 55 °C before being collected by centrifugation with subsequent washes of 50 mM TEAB, 0.2% formic acid, and 50% acetonitrile. Samples were dried and cleaned with GL Science desalting and enrichment ip, GL-Tip. Samples were resuspended in 0.1% trifluoroacetic acid / 2% acetonitrile solution and loaded on to the GL-Tips which were conditioned once with 80% acetonitrile solution in 0.1% trifluoroacetic acid and equilibrated with 0.1% trifluoroacetic acid in 2% acetonitrile solution. The loaded samples were washed once with 0.1% trifluoroacetic acid in 2% acetonitrile solution and finally eluted in 80% acetonitrile solution in 0.1% formic acid.

Peptides were loaded on a 15 cm IonOpticks column which was maintained at 50 °C using a column oven. A Neo Vanquish (Thermo) was directly coupled online with the mass spectrometer (Astral Thermo) and peptides were separated with a binary buffer system of buffer A (0.1% formic acid (FA)) and buffer B (80% acetonitrile plus 0.1% FA), at a flow rate of 400 nl/min. The gradient started at 2% B and increased to 34% in 30 minutes before increasing to 100% within 0.1 minutes and held for 3 minutes prior to returning to 2% and re-equilibrated The mass spectrometer was operated in positive polarity mode with a capillary temperature of 275 °C.

The DIA method consisted of a MS1 scan (*m*/*z* = 380-980) with an AGC target of 5 × 10^6^ and a maximum injection time of 5 ms (R = 240,000). DIA scans were acquired with the Astral detector with an AGC target of 8 × 10^4^. Fragmentation occurred in the HCD cell with a normalised stepped collision energy of 25% and the spectra were recorded in profile mode. 199 non-uniform DIA windows across 380-980 were collected with a maximum injection time of 3ms and a 0.6 s loop control which achieved an average of 5 data points per peak.

MS raw files were processed using DIA-NN (v1.9.2) as described previously (https://doi.org/10.1038/s41467-021-25454-1). Library-free searching was performed with a concatenated databased containing the reviewed human proteime and T. gondii (ME49) proteome. Searching was performed with a maximum of one missed cleavage with MBR enabled and robust LC quantification performed.

The protein intensity matrix was analyzed using R in the integrated development environment RStudio. BioConductor’s Differential Enrichment analysis of Proteomics data (DEP) library served as the backbone of the analysis (https://doi.org/10.1038/nprot.2017.147).. The data was filtered and normalized using variance stabilising normalisation (vsn). Significant outliers were identified using PCA plots for each condition and removed from downstream analysis. The data was imputed with the Bayesian PCA (BPCA) missing value estimation method complemented by QRILC (Quantile Regression Imputation of Left-Censored data). The data was then used to calculate log fold change (logFC) and statistical significance (P-value) for set contrasts of interest. Furthermore, multiple hypothesis correction was conducted using Benjamini-Hochberg correction. The generated adjusted-P values were used for the analysis. The final plots were generated using ggplot library using the above dataset.

## Extended data from Figures

**Extended Figure 1:**
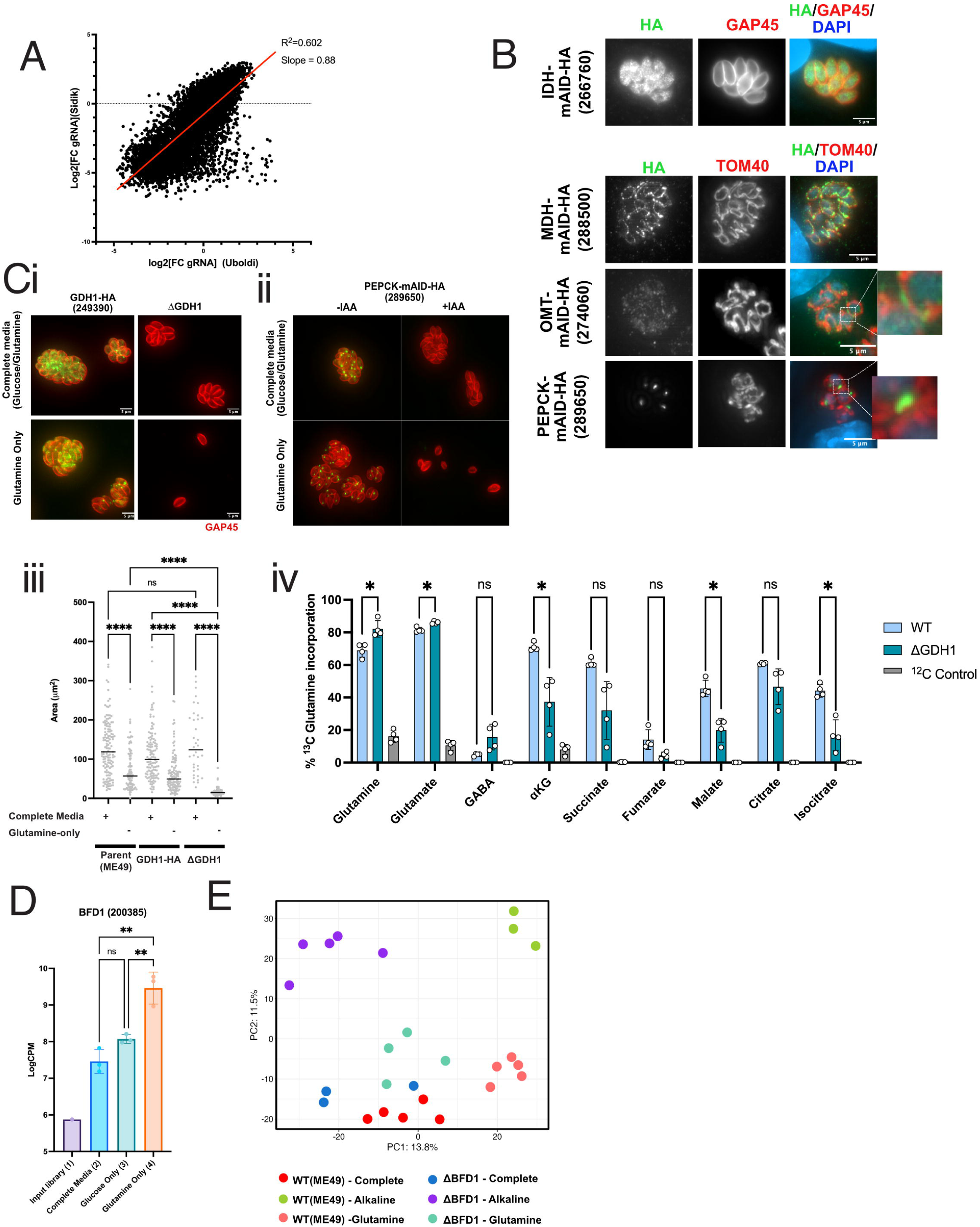
A whole genome CRISPR screen reveals *Toxoplasma* genes involved in utilization of glutamine and glucose. (A) A comparison between CRISPR fitness scores in this study versus Sidik et al ^33^.(B) Localisation of Isocitrate Dehydrogenase (IDH) (266760), Malate Dehydrogenase (MDH) (288500), Oxoglutarate/Malate Transporter (274060) and PEP carboxykinase (PEPCK)(289650) by genetic tagging with HA and mAID. MDH, OMT and PEPCK co-localised with TOM40 mitochondrial marker (Ci) GDH1 (249390) localisation by HA tagging (GDH1-HA) and subsequent knockout ΔGDH1, followed by growth in complete media (Glc^+^/Gln^+^) or in media with glutamine as the carbon source (Glc^+^/Gln^-^). (ii) Localisation and knockdown of PEPCK(289650) by tagging with HA and min-inducible degron (mAID) (PEPCK-mAID-HA). Addition of Indol Acetic Acid (IAA) depletes PEPCK and parasites assessed for growth in either complete media (Glc^+^/Gln^+^) or glutamine only (Glc^-^/Gln^+^). (iii) Quantitation of area of vacuole in mm^2^ of WT(ME49) parent, GDH1-HA and ΔGDH1 in complete (Glc^+^/Gln^+^) and glutamine-only conditions (Glc^-^/Gln^+^). Each data point represents a single parasite vacuole with the bar representing median value. Data set is a single representative experiment to account for growth variation across experiments. Statistical testing done by ordinary one-way ANOVA where ****p<0.0005, ns= not significant. (iv) U-^13^C glutamine labelling (in the presence of pyruvate) in extracellular WT (ME49) parent vs ΔGDH1 mutant lines. The mean +/− SD plotted for n=3 biological repeats. Error bars represent multiple unpaired t-tests where *<0.05 and ns=not significant (D). Abundance of guides targeting BFD1 (200385) as measured by Log (Counts Per Million) (Log2CPM) across conditions of the CRISPR screen as shown in fig 1A. Mean +/− S.D plotted, n=3 biological repeats, One Way ANOVA, where **p< 0.01, ns = not significant. (E) Principle Component Analysis (PCA) global quantitative proteomics of each biological replicate across parasite strains.

**Extended Figure 2:**
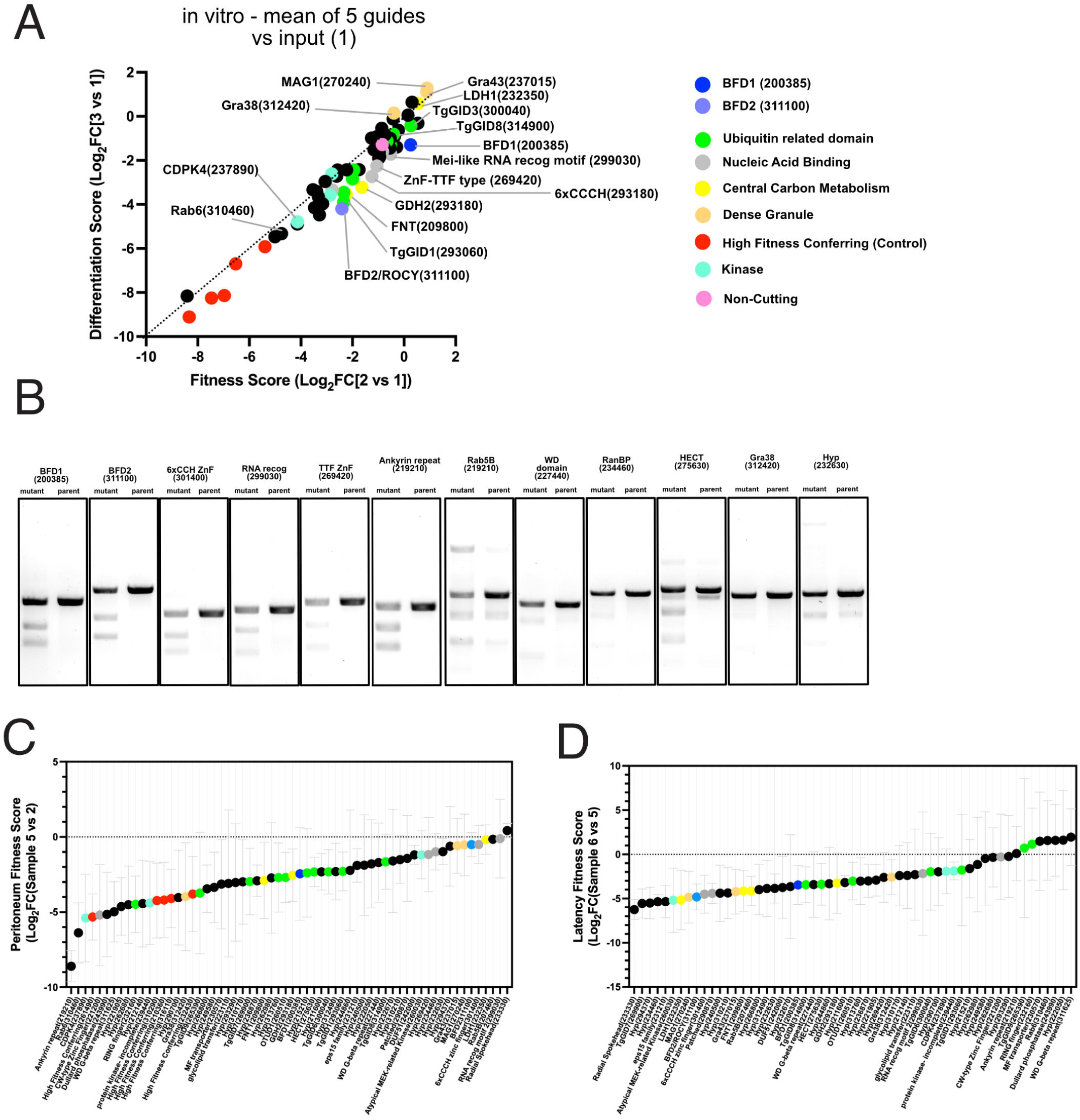
In vitro and in vivo CRISPR screens highlights genes required for differentiation of *Toxoplasma* into latent forms. (A) CRISPR screen results comparing back to input library (Sample 1), showing highly fitness conferring genes that were added to the library (as determined by ^33^). Each datapoint is an individual gene and colour coded according to the key. (B) Surveyor assay to assess gene disruption across mutants generate in Fig 3C as compared to parental line. (C) Waterfall plot of Peritoneum Fitness Scores (Log_2_FC(Sample 5 vs 2). n=3 biological repeats, mean+/− SD shown. (D) Waterfall plot of Latency Fitness Scores (Log_2_FC(Sample 6 vs 5). n=3 biological repeats, mean+/− SD shown.

**Extended Figure 3:**
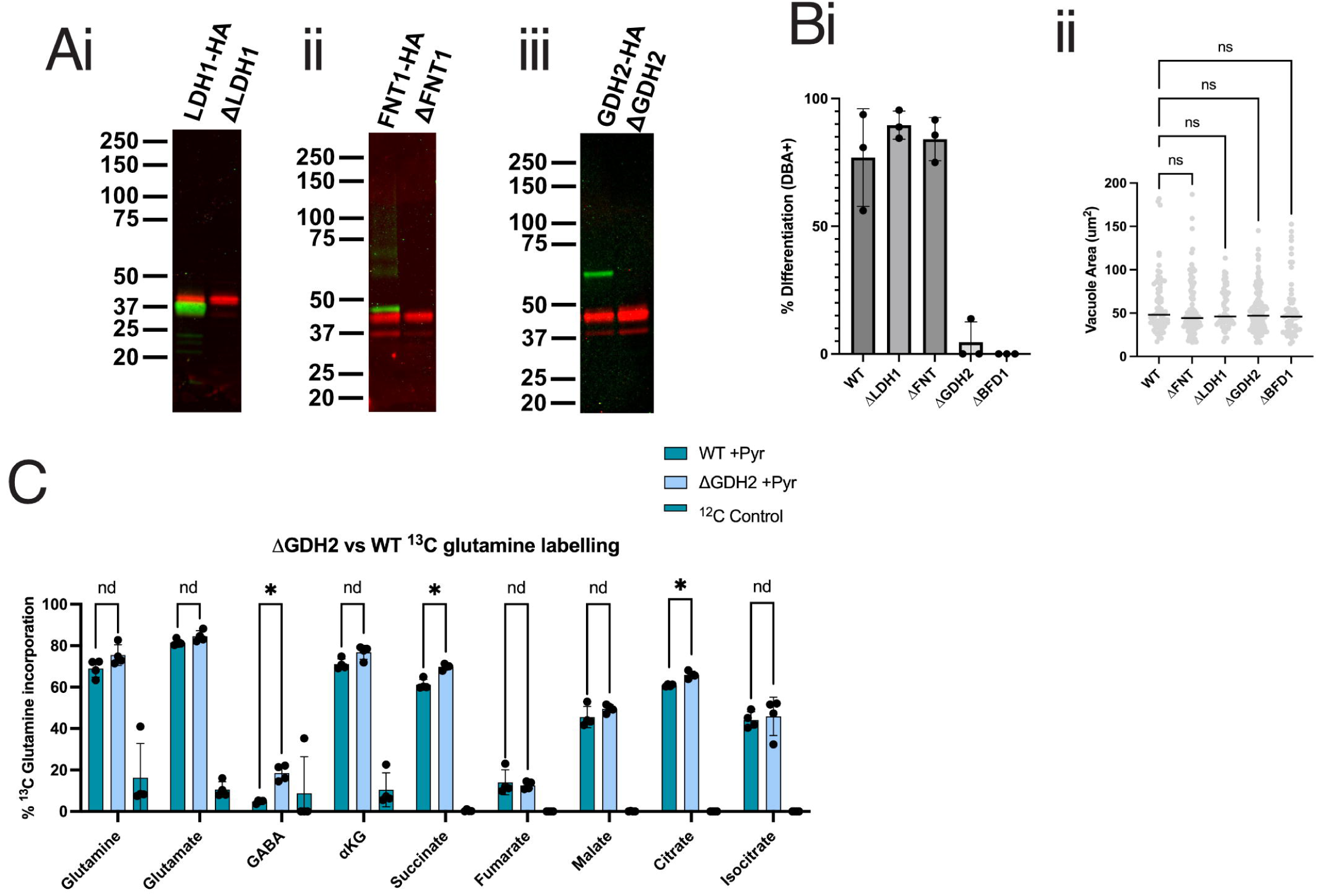
Disruption of *Toxoplasma* genes involved in central carbon metabolism causes defects in differentiation into latent forms. (A) Genetic tagging with HA epitope and knockout of LDH1, GDH2 and FNT1 as confirmed by western blot. HA-Green, aGAP45 – red. (B) Differentiation of WT (parental), ΔLDH1, ΔFNT1, ΔGDH2 and ΔBFD1(control) in HFF, for 48hrs under alkaline stress conditions. (C) Vacuole size of mutants as above, as marked by GAP45 and measured in mm^2^ (D) U^13^C glutamine labelling of extracellular WT vs DGDH2 tachyzoites, showing percentage incorporation of ^13^C label into metabolites of central carbon metabolism. n=4 biological repeats, mean +/− SD shown. Unpaired two tailed t-test used where, *p<0.05 and ns=not significant.

**Extended Figure 4:**
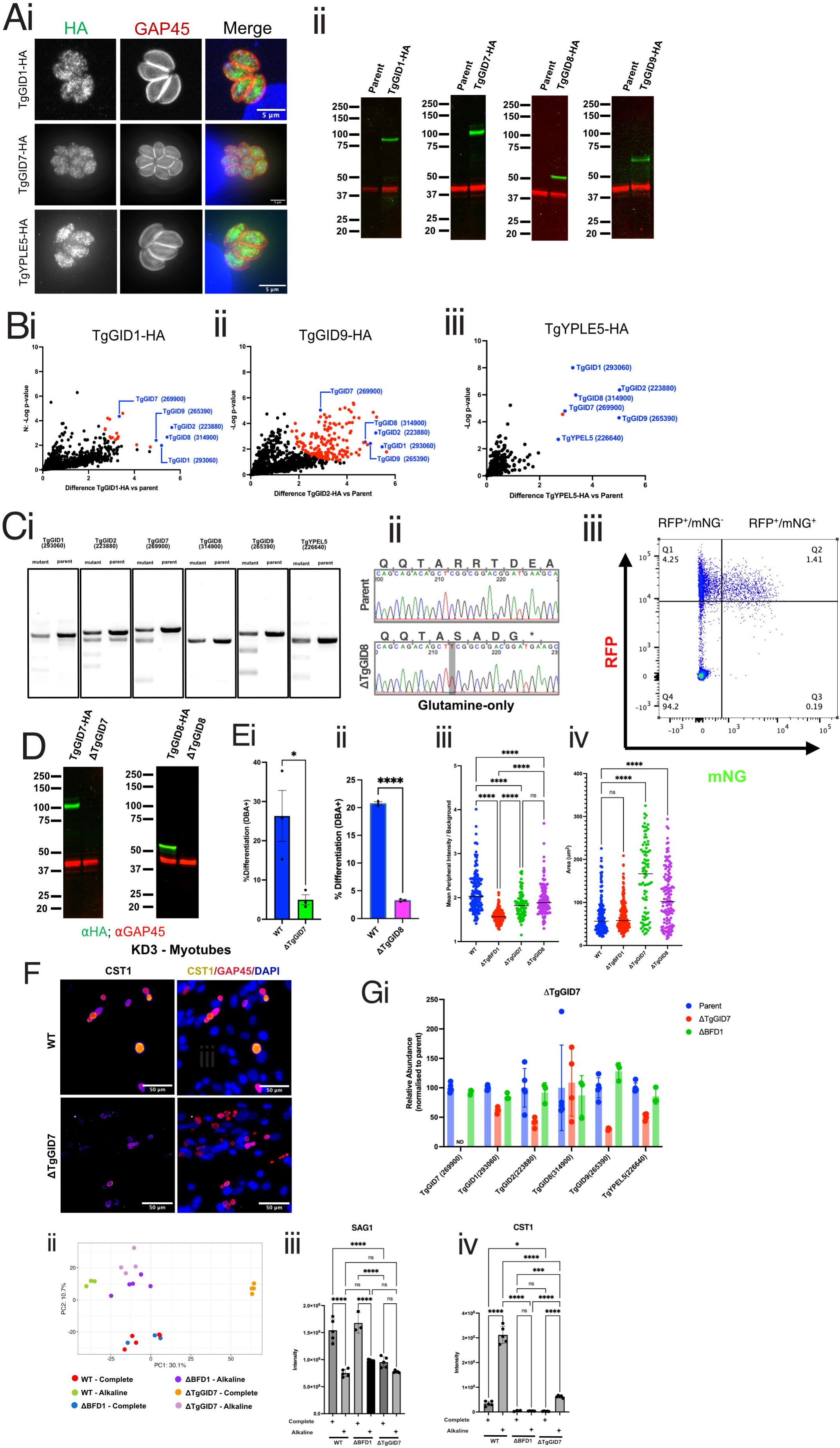
*Toxoplasma* contains a GID E3 ubiquitin ligase complex which is required for differentiation into latent forms. (Ai) Localisation of TgGID1-HA, TgGID8-HA and TgYPEL5-HA. (i) IFA of TgGID1-HA, TgGID7-HA, TgGID8-HA and TgGID9-HA, together with GAP45 (parasite periphery) and DAPI (B) αHA Immunoprecipitation of TgGID1-HA, TgGID8-HA and TgYPEL5-HA as compared to parental control. Statistically significant hits listed in blue and red. Blue = other GID components (C) Genotyping of individual CRISPR mutants generated in the ME49:Cas9 reporter strain ^33^, either by (i) Surveyor assay, as compared to parental control or (ii) sequencing. (iii) Gating strategy for FACS analysis of CRISPR mutants. (D) Western blot confirmation of genetic deletion of TgGID7 and TgGID8. (E) Analysis of differentiation for 48hrs in glutamine-only medium by counting DBA^+^ vacuoles of both (i) ΔTgGID7 vs parent and (ii) ΔTgGID8 vs parent. (iii) Analysis of the mean peripheral intensity of the DBA^+^ signal of ΔTgGID7 and ΔTgGID8 vs WT and ΔBFD1 controls as measured within a 0.1μm thickness around the boundary of the GAP45 signal. Each datapoint represents a single parasite vacuole and bar represents median data point. (iv) Quantification of vacuole size of ΔTgGID7 and ΔTgGID8 as compared to WT parent and ΔBFD1 controls. Area (μm^2^) calculated by GAP45 staining. Graph represents a single representative experiment. One way ANOVA, where ****p<0.0001 and ns = not significant (F). Representative images of differentiation assay in KD3 myoblasts of ΔTgGID7 and WT parental controls. CST1 (fire tone), GAP45 (red), DAPI(blue). (G) Quantitative global proteomic analysis of WT, DBFD1 and ΔTgGID7 grown in complete media or under alkaline stress for 48hrs. (i) Analysis of TgGID subunit abundance in ΔTgGID7 vs WT parent samples from global proteomic analysis. Values normalized to the mean of the abundance of the TgGID subunit in WT parental samples and compared to the levels in ΔBFD1. (ii) Principal component analysis (PCA) of samples. (iii) Protein abundance of the canonical tachyzoite marker SAG1 and (iv) bradyzoite marker CST1 across all conditions. For both (ii) and (iii) n=3-5, mean +/− SD, One way ANOVA, where *p<0.05, **<0.01, ***p<0.001,****p<0.0001 and ns= not significant.

**Extended Figure 5:**
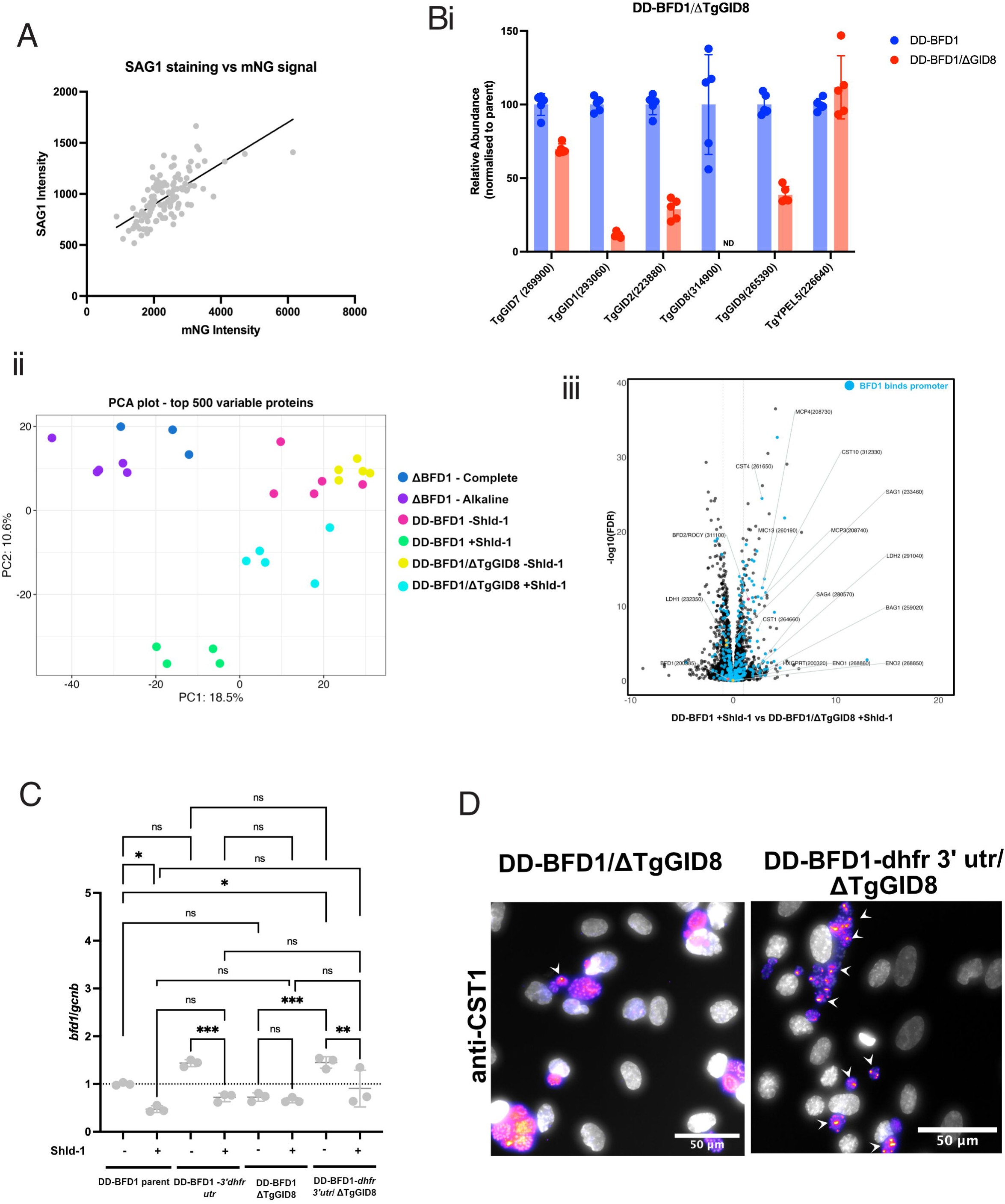
TgGID likely acts through the 3’ utr of *bfd1* to regulate differentiatio. (A) Scatter plot showing the correlation between mNG signal (driven by the *sag1* promoter) and anti-SAG1 staining as measured by quantitative IFAs. (B) Quantitative proteomics of DD-BFD1, DD-BFD1/DGID8 with and without Shld-1 as compared to DBFD1 in complete and alkaline stress conditions. (i) Analysis of TgGID subunit abundance in DD-BFD1/DGID8. Values normalized to the mean of the abundance of the TgGID subunit in DD-BFD1 parental samples (ii) Principal component analysis (PCA) of samples. (iii) Volcano plot comparing of DD-BFD1+Shdl-1 vs DD-BFD1/DTgGID8 +Shld-1. Blue datapoints represents encoding genes whose promoter binds to BFD1 ^12^ and yellow indicates known tachyzoite specific proteins. (C) Quantitative PCR (qPCR) of *bfd1* RNA levels across DD-BFD1 parent and DTgGID8 strains with and without Shld-1 as well as upon swap of the *bfd1* 3’ utr swap to *dhfr* 3’utr. Normalised against *gcnb* similar to Licon et al ^25^. (D) CST1 staining of DD-BFD1/DTgGID8 upon swap of the *bfd1* 3’utr with the *dhfr* 3’ utr. White arrowheads show vacuole with punctate staining of CST1.

## Supplementary Tables

**Table S1: CRISPR screen data**

Tab 1: Whole genome CRISPR screen for growth in glucose-only (Glc^+^/Gln^-^) versus glutamine (Glc^-^/Gln^+^) media from Figure 1.

Tab 2: Sub-library composition of 60 genes. Guides represent first 5 guides as described by Sidik et al ^33^, from Figure 3.

Tab 3: In vitro sub-library CRISPR screen, from Figure 3.

Tab 4:In vivo sub-library CRISPR screen, from Figure 3

**Table S2: Global proteomics of TgGID mutants and carbon source-stimulated differentiation**

**Table S3: IP-MS/MS identification of TgGID complex**

Contains tabs with each immunoprecipitation experiment and a master sheet comparing all data.

**Table S4: Primers, plasmids and resources used in this study**

